# CAG repeat-selective compounds reduce abundance of expanded CAG RNAs in patient cell and murine models of SCAs

**DOI:** 10.1101/2024.08.17.608349

**Authors:** Hannah K. Shorrock, Asmer Aliyeva, Jesus A. Frias, Victoria A. DeMeo, Claudia D. Lennon, Cristina C. DeMeo, Amy K. Mascorro, Sharon Shaughnessy, Hormoz Mazdiyasni, John D. Cleary, Kaalak Reddy, Sweta Vangaveti, Damian S. Shin, J. Andrew Berglund

## Abstract

Spinocerebellar ataxias (SCAs) are a genetically heterogenous group of devastating neurodegenerative conditions for which clinical care currently focuses on managing symptoms. Across these diseases there is an unmet need for therapies that address underlying disease mechanisms. We utilised the shared CAG repeat expansion mutation causative for a large subgroup of SCAs, to develop a novel disease-gene independent and mechanism agnostic small molecule screening approach to identify compounds with therapeutic potential across multiple SCAs. Using this approach, we identified the FDA approved microtubule inhibitor Colchicine and a novel CAG-repeat binding compound that reduce expression of disease associated transcripts across SCA1, 3 and 7 patient derived fibroblast lines and the *Atxn1^154Q/2Q^* SCA1 mouse model in a repeat selective manner. Furthermore, our lead candidate rescues dysregulated alternative splicing in *Atxn1^154Q/2Q^* mice. This work provides the first example of small molecules capable of targeting the underlying mechanism of disease across multiple CAG SCAs.

## Introduction

Microsatellite repeat expansions cause over 50 neurodegenerative, neurological and neuromuscular diseases, including Huntington’s disease (HD), Spinal and bulbar muscular atrophy (SBMA), C9orf72 amyotrophic lateral sclerosis/frontotemporal dementia (C9 ALS/FTD) and myotonic dystrophy types 1 and 2 (DM1, DM2). One of the most frequent class of disease-causing microsatellite expansions are CAG repeat expansions which are the genetic cause of HD, SBMA, Dentatorubral pallidoluysian atrophy (DRPLA), and multiple subtypes of spinocerebellar ataxia (SCA) including types 1, 2, 3, 6, 7, 12 and 17^1^. The SCAs are a genetically heterogenous group of rare neurodegenerative disorders that share the clinical feature of progressive ataxia. This loss of balance and coordination is accompanied by slurred speech, a loss of control of eye movements and reflects dysfunction and degeneration of the cerebellum and interconnected regions of the nervous system including the brainstem. For some SCAs, this dysfunction and degeneration is particularly focused on the Purkinje neurons in the cerebellum, however, for other SCAs, these are less affected^2,3^.

While the precise molecular mechanism(s) responsible for disease pathogenesis remain unclear for CAG SCAs, they share the production of CAG expansion RNAs and the expression of toxic polyglutamine (polyQ) proteins^3,4^. This is the case for SCAs 1, 2, 3, 6, 7 and 17 where the CAG repeat expansion mutation is located within the coding region of *ATXN1*, *ATXN2*, *ATXN3*, *CACAN1A*, *ATXN7*, and *TBP* respectively^1,3,4^. The expanded polyglutamine tracts produced in these diseases alter protein conformation leading to disruption of normal function and protein aggregation^3,4^. However, in the case of SCA12, the CAG repeat expansion is in the 5’UTR of *PPP2R2B* and polyQ proteins have not been reported in this disease^1,3^. As a consequence of the expression of these CAG expansion transcripts and subsequent polyQ proteins, various cellular processes have been implicated in CAG SCAs, including ion channel dysfunction, transcriptional dysregulation, and DNA damage; however, the precise molecular drivers of dysfunction for these pathways remains unclear^3,4^.

Across the SCAs, and indeed CAG expansion diseases, current therapeutic approaches focus primarily on treating symptoms, leaving a large unmet need for therapies that address underlying disease mechanisms in SCAs^2–4^. To target the underlying disease mechanism and the direct consequences of the CAG expansion mutations, several levels of the pathogenic cascade could be targeted: the CAG expansion containing gene, expansion transcripts, or expansion proteins^3^. Alternatively, downstream consequences of this pathogenic cascade can also be targeted^2^. While targeting each level of this pathogenic cascade comes with its own strengths and weaknesses, the further up this cascade that can be successfully targeted, the more likely it is that multiple downstream consequences of disease will be rescued.

While therapeutic strategies targeting each of these levels of the pathogenic cascade are currently in preclinical development for SCAs, one of the more advanced strategies, with clinical trials currently ongoing, is targeting disease specific mRNA transcripts by using antisense oligonucleotides^2–4^. For example, ASOs targeting gene specific sequences reduce levels of the CAG expansion RNAs and the encoded polyglutamine expansion proteins as well as rescuing disease specific phenotypes in SCA1, 2, 3 and 7 mouse models^5–8^. Indeed, a non-allele specific ASO currently in clinical trials for SCA3 (NCT05160558) highlights the promise of this approach^2^. Additionally, short hairpin RNAs and microRNAs that target degradation of expansion mRNAs have also shown therapeutic promise in preclinical studies^9–12^, thereby demonstrating the effectiveness of targeting CAG expansion RNAs in these SCAs.

The majority of these therapeutic approaches have been gene specific, offering hope for one subtype of SCA but not across the larger family of ataxias. Other ASO approaches targeting the CAG repeats directly highlight the potential of shared therapeutics across CAG SCAs. A CAG repeat-targeting ASO has been shown to reduce levels of CAG expansion RNAs and polyglutamine expansion proteins in mouse models of both SCA1 and SCA3^13^ and is currently entering clinical trials for SCA1, 3 and HD (NCT05822908)^2^. Despite this, the suspension of multiple clinical trials for ASOs in HD^2,14^ highlights the challenge of translating ASOs aimed at reducing levels of disease associated RNAs, from preclinical development to approved treatment. Therefore, it is critical to develop alternative therapeutic approaches, including those that could work independently or in combination with ASOs.

Other therapeutic strategies aimed at targeting CAG expansion transcripts, or further upstream in the pathogenic cascade, in a cross-disease manner include RNA cleaving DNAzymes^15^, RNA binding peptides^16,17^, and DNA binding peptides^18^ and small molecules^19,20^. RNA cleaving DNAzymes that lead to degradation of expanded CAG RNA have been shown to reduce polyQ protein levels across models of HD, SCA1, SCA3, SCA7, SCA17 and SBMA^15^. Likewise, the CAG RNA binding peptide BIND, has been shown to reduce levels of polyQ proteins and CAG RNA foci in transfection models of SCA2, as well as rescuing a variety of downstream disease hallmarks in CAG expansion expressing Drosophila and transfection models of SCA2, SCA3 and HD^16,17^. With these approaches, as with the CAG targeting ASO, expansion alleles are preferentially, but not exclusively, targeted due to there being more binding sites for the therapeutic on expansion RNAs than on the equivalent non-expanded RNAs. Furthermore, designer peptides that preferentially bind expanded CAG DNA reduce levels of *HTT* RNA and polyQ aggregates as well as rescue behavior phenotypes in HD models by interfering with transcription elongation^18^. Finally, in both HD and DRPLA mouse models, a CAG expansion DNA binding small molecule, naphthyridine-azaquinolone, has been shown to reduce polyQ aggregates in the striatum by causing CAG repeat contractions^19,20^. Together these studies demonstrate that multiple therapeutic strategies based on a variety of mechanisms can be used to realize therapeutic benefit and rescue multiple levels of the pathogenic cascade across multiple CAG expansion diseases. Despite this, the potential of shared small molecule therapeutics that target the pathogenic source, the CAG repeat expansion, across CAG expansion SCAs remains under-studied with no small molecules currently published to reduce expression of CAG expansion transcripts in SCAs.

To address this therapeutic development gap for this group of rare devastating diseases, we set out to design a small molecule screening system to identify compounds that selectively regulate abundance of CAG RNAs in a disease-independent approach. This innovative disease-gene independent, pathway-agnostic approach has the potential to identify small molecules capable of alleviating the underlying disease mechanisms in CAG expansion SCAs as well as ameliorating downstream cellular consequences. Furthermore, by taking a pathway-agnostic approach, this screening platform has the capability of identifying compounds that regulate expression of expansion RNAs through a variety of mechanisms. Together, this system may therefore identify small molecules with therapeutic potential across multiple CAG expansion SCAs, and potentially across the broader range of CAG microsatellite expansion disorders.

## Results

### Generation and characterization of a HEK293T CAG repeat-selective screening system

Our central hypothesis is that regulating CAG expansion transcript abundance with small molecules will ameliorate downstream cellular consequences and yield therapeutic potential across multiple SCAs. To identify compounds capable of this, we established a CAG expansion-selective screening cell line, based on a similar system we previously reported for DM1^21^. This screening system expresses ATG-(CAG)_60_-Myc-NLuc and ATG-(CAG)_0_-Myc-FLuc transcripts, each with a unique qPCR probe binding site 3’ of the repeat tract to distinguish the two transcripts. This permits the sensitive ratiometric measurement of polyglutamine-nanoluciferase (polyQ-NLuc) relative to firefly luciferase (FLuc) using a dual luciferase assay, and measurement of r(CAG)_60_ relative to r(CAG)_0_ using a multiplex RT-qPCR with one common primer pair and fluorescent probes against the unique sequences (Fig 1A). These constructs do not contain genetic context from any SCA disease causing genes. The constructs were randomly integrated into the HEK293T cell genome using the PiggyBAC Transposon system, cells were selected for puromycin resistance, single cell sorted, and screened for expression of both integrands. Low, medium, and high expressing clones (15, 10, 37 respectively) were identified, with expression of both integrands confirmed via qPCR (Fig 1B), dual luciferase assay (Fig 1C) and protein blotting using antibodies against the myc epitope tag, FLuc, NLuc and polyQ (Fig 1D).

**Figure 1.**
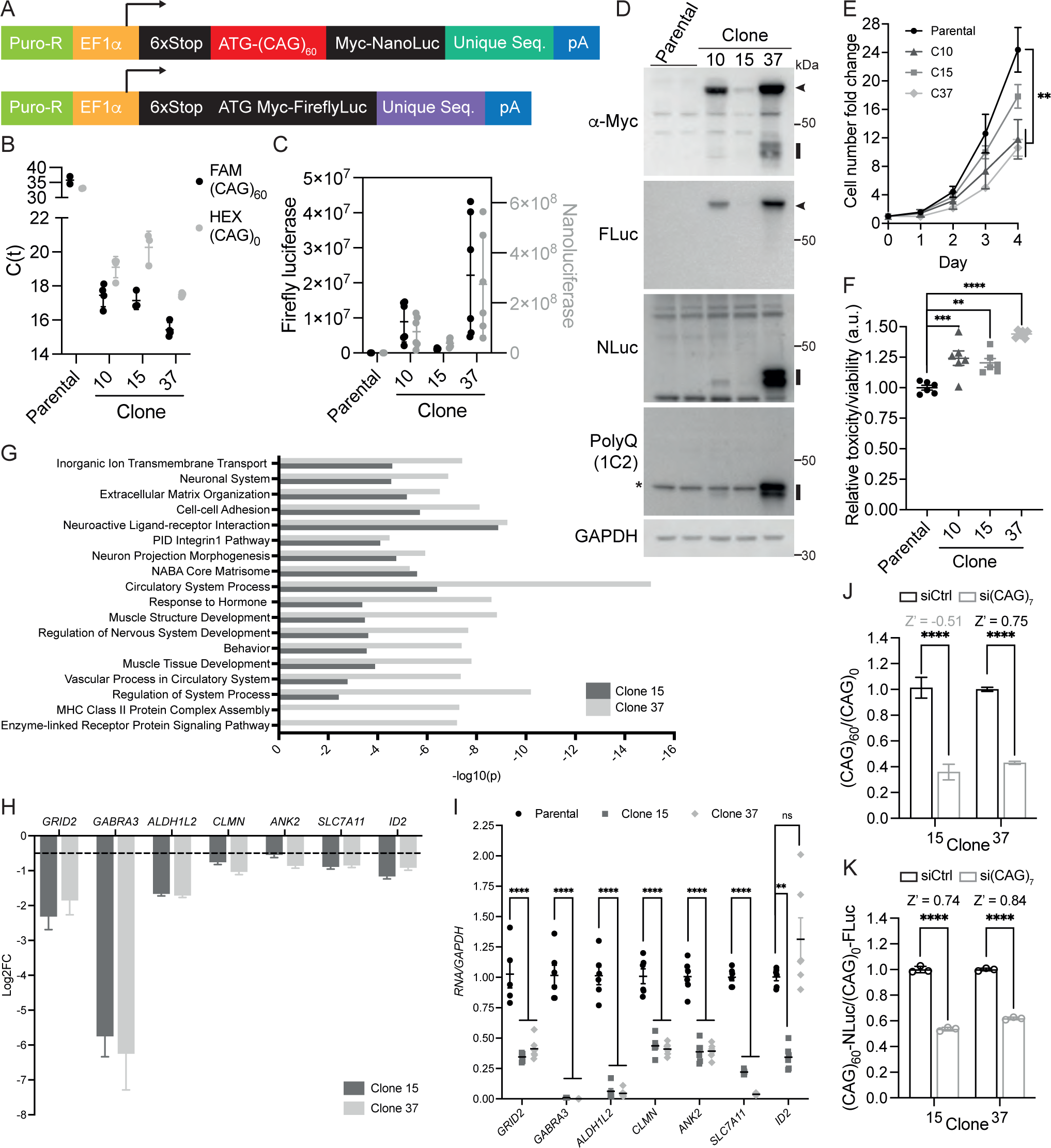
Establishing a disease-independent CAG repeat-selective screening cell system. **A** Schematic of constructs integrated into HEK293T genome to generate ATG-(CAG)_60_-Myc-NLuc/ATG-(CAG)_0_-Myc-Fluc cell line; Puro-R: puromycin resistance gene; EF1α: human EF1α promoter; 6xStop: 2 stop cassettes in each reading frame. **B** Expression of (CAG)_60_ and (CAG)_0_ in parental HEK293T cells and clones 10, 15 and 37 detected measured using FAM and HEX labelled probes respectively, in a multiplex qPCR; C(t): cycle threshold. **C** Dual luciferase assay for protein expression for polyQ-NLuc and FLuc in parental HEK293T cells and clones 10, 15 and 37. **D** Protein blotting for polyQ-Myc-NLuc and Myc-FLuc in parental HEK293T cells and clones 10, 15 and 37; Arrow heads indicate FLuc, black bars indicate polyQ-NLuc,*-TATA binding protein. **E** Cell number fold change for parental HEK293T cells and clones, 10, 15 and 37; n≥4 per cell line per timepoint, mean ± SEM, ** *P*<0.01, one-way ANOVA with Tukey’s multiple comparisons test for day 4. **F** Relative cell toxicity normalized to cell viability for parental HEK293T cells and clones 10, 15 and 37; n=6, mean ± SEM; ** *P*<0.01, *** *P*<0.001, **** *P*<0.0001, one-way ANOVA with Tukey’s multiple comparisons test. **G** Enrichment of summary gene ontology terms identified using Metascape. **H, I** RNA-Seq log_2_fold change (log_2_FC) (**H**) and qPCR validation (**I**) for seven genes significantly differentially expressed in a consistent direction in clone 15 and 37 compared to parental cell lines, and in CAG SCA mouse models. **H** Log_2_FC>0.5 (indicated by dashed line), *P*adj<0.05, n=3. **I** n=6, mean±SEM, one-way ANOVA with Tukey’s multiple comparisons test, ns – not significant, ** *P*<0.01, **** *P*<0.0001. **J, K** CAG targeting siRNA [si(CAG)_7_] selectively reduces levels of CAG expansion transcripts (**J**) and polyQ proteins (**K**). **J** clone 15 n≥5, clone 37 n=3, mean±SEM; **K** n=3, mean±SD. **** p<0.0001, two-tailed unpaired t-test; Z’ scores between 0.5 and 1 indicate that the assay is suitable for small molecule screening^31^.

Upon characterization of the clonal cell lines, it was noted that all clonal lines grew slower than parental HEK293T cells, with clone 10 and clone 37 showing a 12.6% (*P*=0.0047) and 13.8% (*P*=0.0022) reduction in cell number fold change (growth rate) compared to parental four days after plating, respectively (Fig 1E). This reduction in growth rate was accompanied by a 44.0% (*P*<0.0001) increase in relative cell toxicity for clone 37 compared to parental HEK293T cells. For clone 10 there was a 24.2% (*P=*0.0007) and for clone 15 a 20.4% (*P*=0.004) increase in cell toxicity compared to parental, thereby revealing a dose effect of increasing CAG repeat load with increasing cell toxicity. High and low expressing clones (clone 37 and clone 15) were selected for further characterization by RNA sequencing (RNA-Seq) analysis (Table S1). Compared to parental, both clonal lines demonstrated a greater number of downregulated genes compared to upregulated genes (Fig S1A, B) consistent with differential gene expression patterns seen in mouse models of CAG SCAs^22–27^. The identified differential gene expression changes occurred in pathways reported to be affected in CAG SCA mouse models such as ion transmembrane transport, neuron projection morphogenesis and neuroactive ligand-receptor interaction (Fig 1G). Multiple genes previously reported to show differential gene expression in a variety of CAG SCA mouse model cerebellum^23,25,28–30^ were identified as being differentially expressed in both clone 15 and 37 (Fig S1C). Several of these genes showed changes in transcript expression in the same direction as reported in CAG SCA mouse models and were validated via qPCR. These included *GRID2* and *GABRA3* which encode ion channel subunits, and *CLMN* and *ANK2* which encode proteins that are associated with the actin cytoskeleton (Fig 1H, I, Fig S1D-F). Together, these data demonstrate that CAG expansion containing clonal cell lines display phenotypic and molecular hallmarks consistent with mouse models of CAG SCAs.

To enable small molecule screening in this system, we sought to identify a suitable positive control from the literature. Because no compounds have been published to selectively reduce CAG expansion transcript levels, we utilized a CAG targeting siRNA [si(CAG)_7_], based on the CAG targeting ASO studied in the context of SCA1 and SCA3^13^, as a positive control with a scrambled siRNA (siCtrl) as a negative control. Clone 15 showed a 65.4% selective reduction in (CAG)_60_ relative to (CAG)_0_ and a 46.4% reduction in polyQ-NLuc relative to FLuc (both *P*<0.0001) on si(CAG)_7_ treatment compared to scrambled siRNA. For clone 37, there was a 57.0% selective reduction of (CAG)_60_ and a 38.1% selective reduction of polyQ-NLuc (both *P*<0.0001) relative to the no-repeat controls for si(CAG)_7_ treatment compared to scrambled siRNA (Fig 1J, K). The Z’ scores between 0.5 and 1 (Fig 1J, K) indicate that this assay is suitable for screening^31^ at the protein and RNA levels in clone 37 and at the protein level in clone 15.

### The FDA approved compound Colchicine regulates CAG expansion RNA levels

To ensure we had an appropriate comparison for screening a small molecule library prepared in DMSO, we sought a small molecule positive control that could regulate expression of CAG expansion transcripts. Based on its ability to regulate expression of CUG expansion RNAs^21^, we tested and identified the microtubule inhibitor colchicine as a DMSO-soluble small molecule regulator of CAG expansion transcript abundance (Fig 2). In clone 37, a 24 hour treatment of 0.1nM colchicine resulted in a 46.9% selective reduction (*P*<0.0001) of (CAG)_60_ levels compared to DMSO treatment, with a Z’ score of 0.6 (Fig 2A) indicating an assay suitable for small molecule screening^31^. In clone 15, 0.1nM colchicine treatment for 48 hours resulted in a selective reduction of CAG expansion transcript levels by 57.1% (*P*=0.0159; Fig 2B). In both clonal cell lines and parental HEK293T cells, colchicine did not induce a reduction of cell viability within and beyond the effective dose range (Fig 2C).

**Figure 2.**
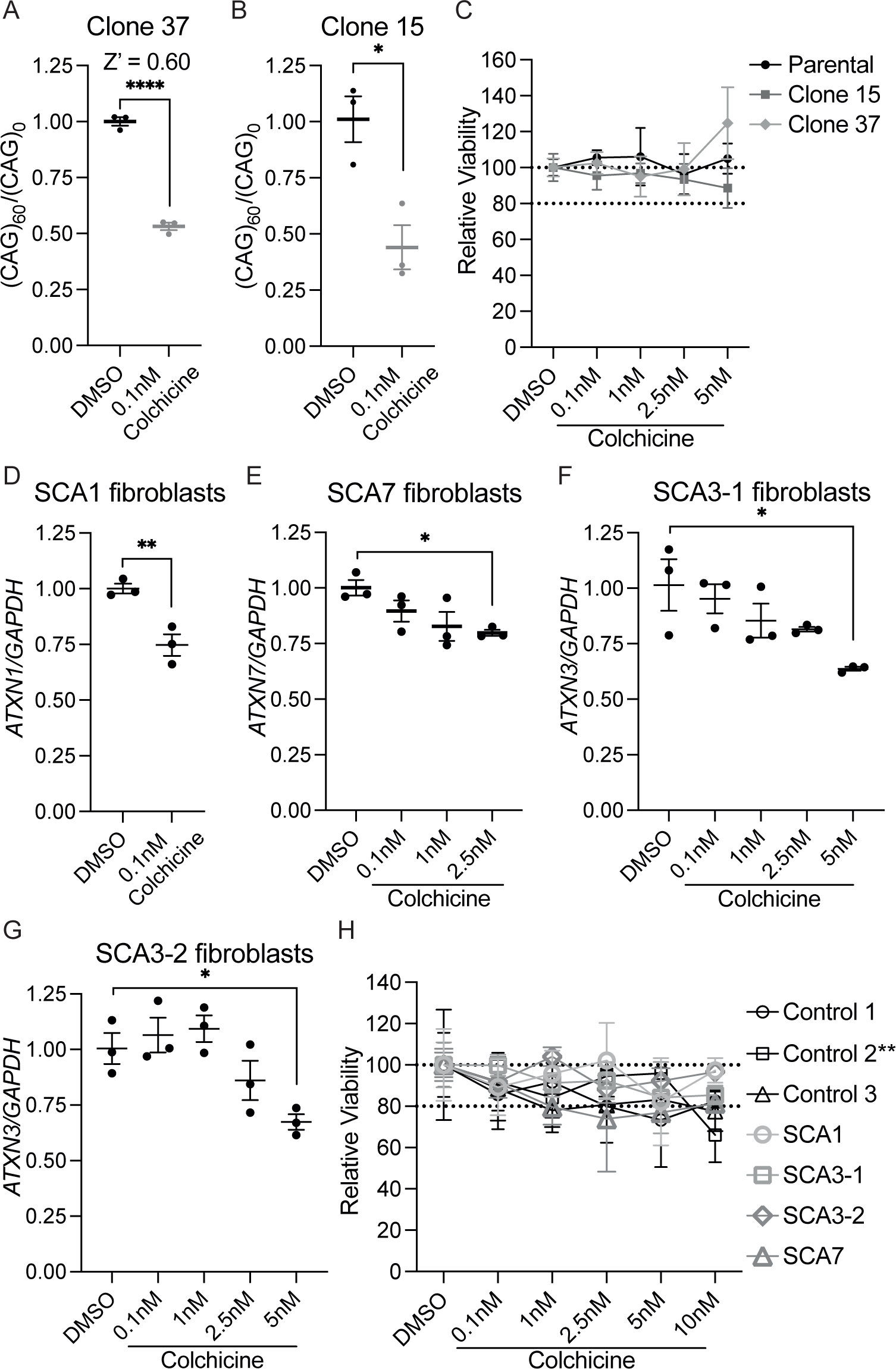
Colchicine regulates expression of CAG expansion transcripts. **A, B** Clone 37 (**A**) and clone 15 (**B**) show a reduction in (CAG)_60_ levels relative to (CAG)_0_ following 0.1nM colchicine treatment; n=3, mean±SEM, two-tailed unpaired t-test, * *P*<0.05, **** *P*<0.0001; Z’ scores between 0.5 and 1 indicate that the assay is suitable for small molecule screening^31^. **C** Relative cell viability of parental, clone 15 and clone 37 cells following 24 hours colchicine treatment compared to DMSO; dotted lines indicate 100% and 80% cell viability, n≥3, mean±SD. **D-G** Colchicine reduces expression of *ATXN1* transcripts in SCA1 patient-derived fibroblasts (**D**), *ATXN7* in SCA7 fibroblasts (**E**), and *ATXN3* in two SCA3 fibroblast lines (**F, G**); n=3, mean±SEM; unpaired two-tailed t-test (**D**), one-way ANOVA with Tukey’s multiple comparisons test (**E-G**), * *P*<0.05, ** *P*<0.01. **H** Relative cell viability of three control, one SCA1, two SCA3 and one SCA7 fibroblast lines following 48 hours treatment with colchicine; dotted lines indicate 100% and 80% relative viability; n=4, mean±SD, two-way ANOVA with Tukey’s multiple comparisons test; Control 2 shows a significant reduction in viability at 10nM compared to DMSO: ** *P*<0.01; all other cell lines show no significant difference between DMSO and 10nM.

To further validate the use of colchicine as a positive control small molecule in CAG SCAs, we treated SCA1, SCA3 and SCA7 patient derived fibroblast lines with colchicine for 48 hours. In an SCA1 patient derived cell line, 0.1nM colchicine reduced expression of *ATXN1* by 25.3% (*P*=0.009; Fig 2D). In SCA3 and SCA7 no effect was seen at 0.1nM for 48 hours so a wider dose range was tested (Fig 2E-F). In an SCA7 patient-derived fibroblast line, 2.5nM colchicine led to a 20.2% reduction in *ATXN7* expression levels (*P*=0.049; Fig 2E). Two different SCA3 patient derived cell lines showed a reduction of *ATXN3* expression following treatment with 5nM colchicine: in the SCA3-1 line, this was a reduction by 37.7% (*P*=0.021; Fig 2F), and in SCA3-2 a reduction of *ATXN3* expression by 33.1% (*P*=0.042; Fig 2G). There was, however, no effect in three control fibroblast lines of 0.1nM colchicine on *ATXN1* expression (Fig S2A) or of 5nM colchicine on *ATXN3* expression (Fig S2C) and two out of three control fibroblast lines showed a significant increase in expression of *ATXN7* following treatment with 2.5nM colchicine (Control 2: 59.5% increase, *P*=0.013; Control 3: 81.5% increase, *P*=0.0002; Fig S2B). These data demonstrate that colchicine regulates CAG transcript levels in a repeat length dependent manner and suggests differences in the regulation of control allele expression for *ATXN7* versus *ATXN1* and *ATXN3*.

Although colchicine treatment was well tolerated in clone 15 and 37 (Fig 2C), an overall trend of reduced cell viability was seen for colchicine treatment in fibroblasts but only control 2 showed a significant reduction in cell viability at the highest dose tested (10nM, 33.9% reduction, *P*=0.002; Fig 2H). Finally, despite promising data of colchicine on RNA expression, in both clone 15 and clone 37, colchicine treatment failed to selectively reduce poly-NLuc expression levels (Fig S2D, E), perhaps suggesting that sufficient expansion RNA is still being transcribed and exported to the cytoplasm to maintain consistent levels of polyQ proteins. Together, these data provide proof-of-concept that small molecules can regulate CAG expansion transcripts in a repeat length dependent manner across multiple CAG expansion diseases.

### A CAG repeat selective screen identifies a pyrazole-based compound that regulates CAG expansion RNA levels

To maximize the potential of identifying different structural classes of compounds with the potential to regulate CAG transcript abundance through different mechanisms, we performed a screen of 1584 structurally diverse compounds from the National Cancer Institute Developmental Therapeutics Program (NCI DTP) Diversity Set VI. Compounds were screened at 1μM in 0.01% DMSO in clone 37 for 24 hours, utilizing both colchicine and si(CAG)_7_ as positive controls and scrambled siRNA and DMSO as negative controls (Fig 3). Across the screen, 0.1nM colchicine showed a 19.6% reduction in (CAG)_60_ relative to (CAG)_0_ (*P*<0.0001; Fig 3A) and CAG targeting siRNA selectively reduced (CAG)_60_ abundance by 24.5% (*P*<0.0001; Fig S3A). Control reverse transcriptase reactions were used to confirm that the RT-qPCR screening assay behaved as expected (Fig S3B). The screen of 1584 Diversity set VI compounds identified 72 compounds that selectively reduced (CAG)_60_ expression beyond the 90% confidence interval (CI) of the mean for both positive controls (lower 90% CI of mean: 0.1nM colchicine=77.6%, si(CAG)_7_=70.6%) reflecting a hit rate of 4.54% (Fig 3B).

**Figure 3.**
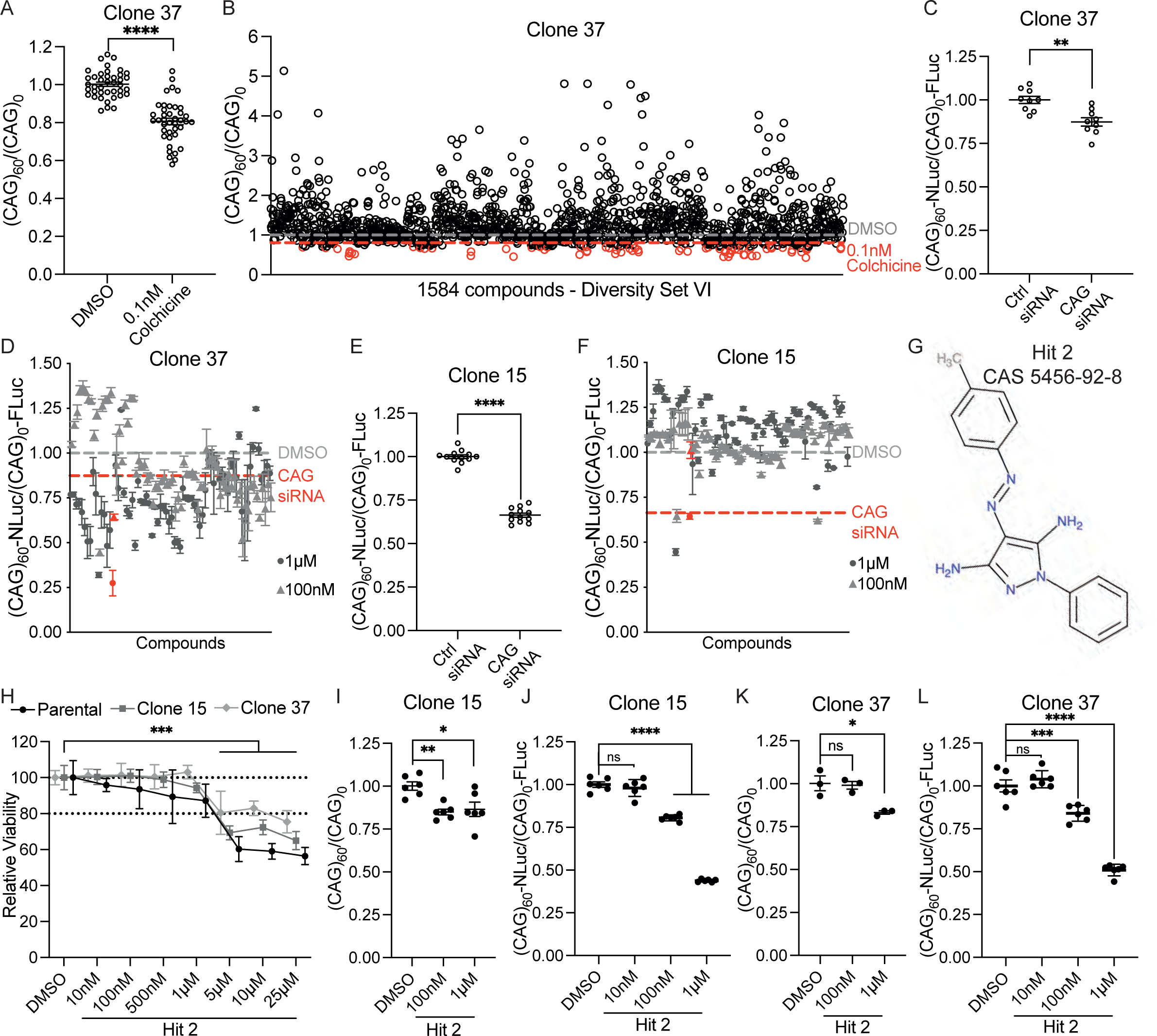
A screen of 1584 structurally diverse compounds identifies a pyrazole-based small molecule as a selective regulator of CAG expansion transcript abundance. **A** 24 hour treatment of clone 37 with 0.1nM colchicine selectively reduces (CAG)_60_ expression (normalized to (CAG)_0_); n=40 (n=2 per screening plate), mean±SEM, unpaired two-tailed t-test, **** *P*<0.0001. **B** Diversity set VI screen results for each individual compound treatment (1µM in 0.01% DMSO) on (CAG)_60_ normalized to (CAG)_0_ for clone 37. Dashed grey line indicates DMSO average and dashed red line indicates colchicine average from Fig 3A. Compounds that passed screening threshold are shown in red. **C, E** 24 hour treatment of clone 37 (**C**) and clone 15 (**E**) with si(CAG)_7_ selectively reduces polyQ-NLuc expression (normalized to FLuc); n=12 (n=2 per screening plate), mean±SEM, unpaired two-tailed t-test, ** *P*<0.01, **** *P*<0.0001. **D, F** Screen results for each individual compound treatment in clone 37 (**D**) and clone 15 (**F**) on polyQ-NLuc normalized to FLuc, for the 72 hits identified in Fig 3B. Dashed grey line indicates DMSO average and dashed red line indicates si(CAG)_7_ average from Fig 3C or Fig 3E for clone 37 and clone 15, respectively. The three compounds that pass threshold in **F** are referred to, from left to right, as Hit 1, 2 and 3. Lead candidate Hit 2 is shown in red; n=3, mean±SD. **G** Chemical structure and CAS number for Hit 2; IUPAC name: 4-[(4-methylphenyl)diazenyl]-1-phenylpyrazole-3,5-diamine; molecular formula: C_16_H_16_N_6_; molecular weight: 292.34 g/mol. **H** Relative cell viability of parental, clone 15 and clone 37 cells following 24 hours Hit 2 treatment normalized to DMSO per cell line; dotted lines indicate 100% and 80% cell viability, clone 15 and clone 37 n=6, parental n≥3, mean±SD, one-way ANOVA with Tukey’s multiple comparison test, *** *P*>0.001. **I, K** 24 hour treatment of clone 15 (**I**) and clone 37 (**K**) with Hit 2 in the tolerated dose range, reduces (CAG)_60_ expression (normalized to (CAG)_0_); clone 15 n=6, clone 37 n=3; mean±SEM, one-way ANOVA with Tukey’s multiple comparisons test, ns – not significant, * *P*<0.05, ** *P*<0.01. **J, L** 24 hour treatment of clone 15 (**J**) and clone 37 (**L**) with Hit 2 in the tolerated dose range, selectively reduces polyQ-NLuc expression (normalized to FLuc); n=6, mean±SD, one-way ANOVA with Tukey’s multiple comparisons test, ns – not significant, *** *P*<0.001, **** *P*<0.0001.

To ensure compounds identified in this initial RNA expression screen (Fig 3B) also selectively reduced polyQ protein levels and were acting independent of construct integration sites, the 72 compounds were rescreened at 1µM and 100nM in both clone 15 and clone 37 for selective reduction of polyQ levels and in clone 15 for selective reduction of CAG RNA levels. For clone 37, the CAG targeting siRNA selectively reduced polyQ-NLuc expression by 12.6% (*P*=0.0011; Fig 3C) with 65 out of 72 compounds selectively reducing polyQ-NLuc expression below this threshold for at least one dose (Fig 3D). In clone 15, si(CAG)_7_ selectively reduced expression of (CAG)_60_ by 37.6% (*P*<0.0001, Fig S3C) and as with the RNA expression screen in clone 37, control reverse transcriptase reactions were used to confirm that the RT-qPCR screening assay behaved as expected in clone 15 (Fig S3D). Colchicine selectively reduced (CAG)_60_ by 20.1% (*P*<0.0001, Fig S3E) in clone 15 with 15 of 72 compounds selectively reducing (CAG)_60_ expression below this threshold for at least one dose (Fig S3F). In clone 15, si(CAG)_7_ selectively reduced expression of polyQ-NLuc by 33.6% (*P*<0.0001; Fig 3E) with three compounds (Hits 1, 2 and 3) selectively reducing expression of polyQ-NLuc below this threshold (Fig 3F); all three compounds also passed the threshold for reduction of (CAG)_60_ expression in clone 15 (Fig S3F). Of these three Hit compounds, Hit 2 showed the greatest selective reduction of poly-NLuc expression in clone 37 (72.6%; Fig 3D) and was the third best performing compound in the clone 37 RNA expression screen (selective reduction of (CAG)_60_ by 54.5%; Fig 3B) out performing Hit 1 and 3 by >20%. Therefore, Hit 2 was selected for further investigation as a regulator of CAG expansion expression.

Hit 2 (IUPAC name: 4-[(4-methylphenyl)diazenyl]-1-phenylpyrazole-3,5-diamine; CAS number: 5456-92-8) is a pyrazole based compound (Fig 3G) soluble in DMSO and methanol, but not in PBS or water. Hit 2 has limited published data for cell-based assays and so we initially set out to identify a dose range that did not negatively impact cell viability in HEK293T cells. In parental HEK293T cells and clones 15 and 37, Hit 2 was well tolerated up to 1µM but at 5µM, 10µM and 25µM, Hit 2 significantly reduced cell viability across all three lines. In clone 37 there was a 17% or greater reduction in cell viability at doses greater than 1µM, for clone 15 this was a >27% reduction in viability and >39% reduction in viability for parental HEK293T cells compared to DMSO treatment for each cell line (*P*>0.001; Fig 3H). Beyond 25µM, Hit 2 crashed out of solution in cell culture media and formed crystals. Within the tolerated dose range, clone 15 showed a 15.2% (*P*=0.0066) and a 13.6% (*P*=0.0142) selective reduction in (CAG)_60_ expression following 24 hour treatment with 100nM and 1µM Hit 2, respectively (Fig 3I). This corresponded to a 19.5% and 56.2% selective reduction (both *P*<0.0001) in polyQ-NLuc expression at 100nM and 1µM of Hit 2, respectively (Fig 3J). Similarly, in clone 37, at 100nM, while Hit 2 did not significantly reduce expression of (CAG)_60_, there was a 16.0% reduction in polyQ-NLuc (*P*=0.0006) and at 1µM there was a 17% selective reduction (*P*=0.0136) in (CAG)_60_ and a 49.1% reduction in polyQ-NLuc (*P*<0.0001; Fig 3K, L). Together these data demonstrate that Hit 2 is a novel compound capable of regulating expression of CAG expansion transcripts and polyQ proteins. The larger reduction in polyQ protein levels suggests that the mechanism of action has a greater impact on inhibiting translation than on reducing CAG expansion transcript abundance.

### Colchicine and lead candidate compound Hit 2 regulate CAG expansion RNA abundance through distinct mechanisms

We previously demonstrated that the mechanism of action of the microtubule inhibitor colchicine in DM1 is dependent on CUG repeats being integrated into the cell genome rather than being expressed ectopically via plasmid overexpression^21^. To test if this was consistent for the mechanism of action of colchicine on CAG RNA expression, we performed transient co-transfections of the ATG-(CAG)_60_-Myc-NLuc and ATG-(CAG)_0_-Myc-FLuc constructs used to generate clone 15 and 37 screening cell lines. Following transient co-transfection, our CAG targeting siRNA positive control selectively reduced (CAG)_60_ expression relative to (CAG)_0_ expression by 52.8% (*P*<0.0001; Fig 4A) consistent with the effect seen in clone 15 and clone 37 (Fig 1J). Treatment with colchicine at 0.1nM, 1nM and 5nM did not affect expression of (CAG)_60_ in the co-transfection system (Fig 4B). These data are consistent with the mechanism of action through which colchicine regulates expression of CUG expansion RNAs^21^. In contrast to colchicine, treatment of the co-transfection system with Hit 2 at 100nM and 1µM reduced (CAG)_60_ expression relative to (CAG)_0_ expression by 26.8% (*P*=0.0175) and 28.8% (*P*=0.0110), respectively (Fig 4C). This plateauing effect of Hit 2 treatment on (CAG)_60_ expression is consistent with the effect of Hit 2 in clone 15 (Fig 3I). Similar to the dose dependent effect of Hit 2 on polyQ-NLuc expression in both clone 15 and clone 37 (Fig 3J, L), in the co-transfection system Hit 2 selectively reduced polyQ-NLuc expression by 58.1% at 10nM, 70.5% at 100nM, and 90% at 1µM (all *P*<0.0001; Fig 4D). Together these data demonstrate that Hit 2 and colchicine regulate expression of CAG expansion RNAs through distinct mechanisms and confirm that Hit 2 has a greater effect on selectively reducing expression of polyQ-NLuc proteins compared to (CAG)_60_ RNAs.

**Figure 4.**
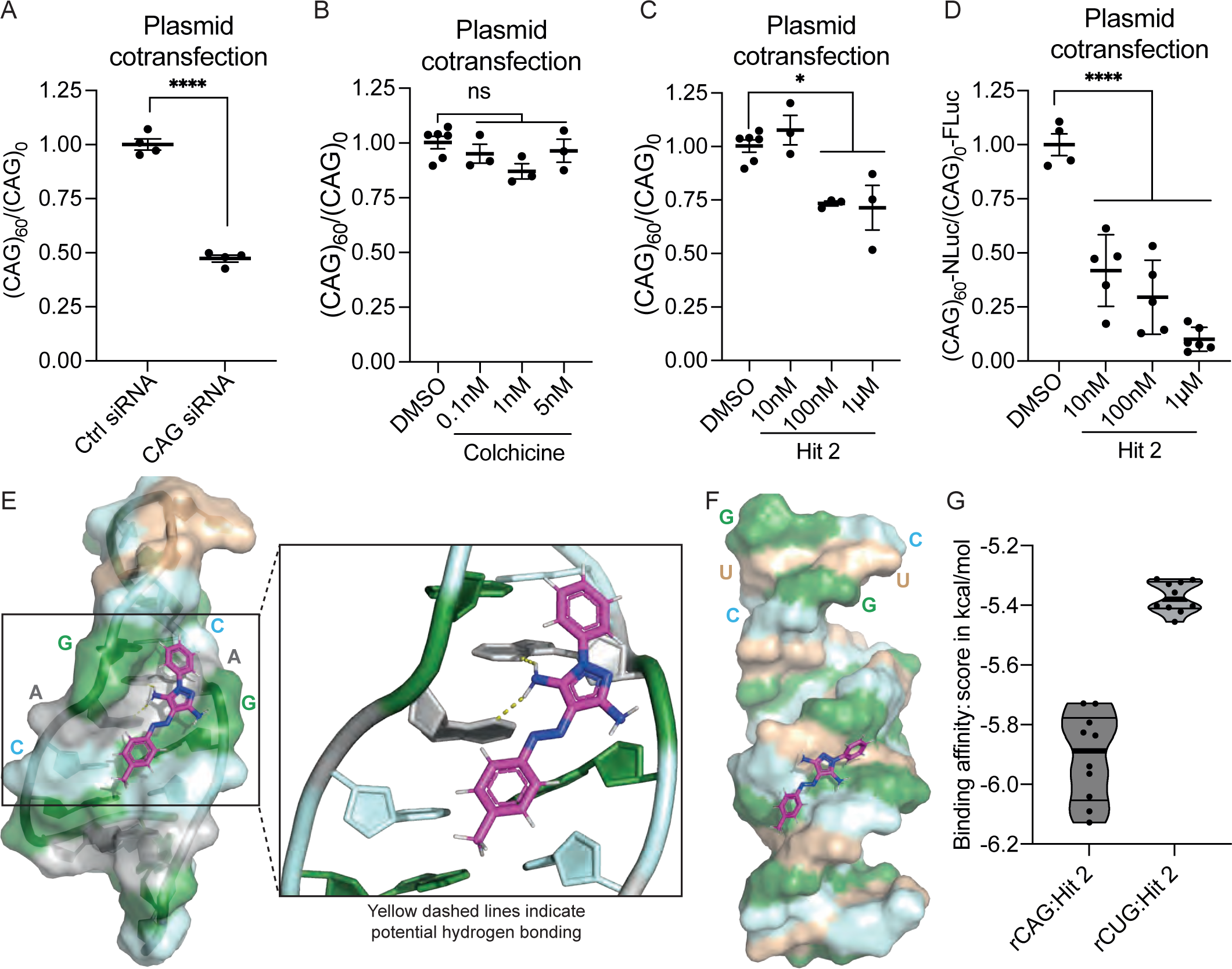
Hit 2 and colchicine regulate CAG expansion transcript abundance through distinct mechanisms. **A-C** 24 hour treatment of ATG-(CAG)60-Myc-NLuc and ATG-(CAG)0-Myc-FLuc co-transfection in HEK293T cells with si(CAG)_7_ (**A**), or Hit 2 (**C**) in the tolerated dose range, reduce (CAG)_60_ expression (normalized to (CAG)_0_), but treatment with colchicine (**B**) in the tolerated dose range does not affect (CAG)_60_ expression. **A** n=4, unpaired two-tailed t-test; **B, C** DMSO n=6, colchicine and Hit 2 n=3, one-way ANOVA with Tukey’s multiple comparisons test. mean±SEM, ns – not significant, * *P*<0.05, **** *P*<0.0001. **D** 24 hour treatment of ATG-(CAG)60-Myc-NLuc and ATG-(CAG)0-Myc-FLuc co-transfection in HEK293T cells with Hit 2 in the tolerated dose range, selectively reduces polyQ-NLuc expression (normalized to FLuc); n≥4, mean±SD, one-way ANOVA with Tukey’s multiple comparisons test, **** *P*<0.0001. **E, F** Computational modelling of a CAG repeat RNA containing a CAG helical region and a UUCG loop (from PBD ID: 7D12) (**E**) and a CUG RNA duplex (from PBD ID: 1ZEV) (**F**) with Hit 2 binding to the minor groove. Magnified site view in **E** highlights hydrogen bonding (dotted yellow lines) between Hit 2 amine group and AA stacked mismatch. **G** Docking scores representing binding affinity indicate Hit 2 binds more efficiently to CAG repeat RNA than CUG repeat RNA; thick line indicates median.

To understand more about the potential mechanism of action of Hit 2, we performed molecular docking studies to investigate if Hit 2 has the potential to bind expansion RNAs. *In silico* molecular modelling and docking predicts that Hit 2 preferentially binds to the minor groove of CAG repeat RNA. This binding allows for hydrogen bonds to form between one of the amine groups on Hit 2 and the AA stacked mismatch in the CAG RNA helix (Fig 4E). Interestingly, molecular docking predicts that Hit 2 also prefers the minor groove when interacting with a CUG RNA duplex (Fig 4F), however, this is a less favorable interaction with a weaker binding affinity (–5.38 kcal/mol) than for Hit 2 binding to CAG repeat RNA (–5.89 kcal/mol; Fig 4G). Together these molecular docking studies suggest that Hit 2 adopts a conformation that is suitable for binding to the minor groove of CAG repeat RNA where the interaction is stabilized by hydrogen bonds between Hit 2 and the AA stack. Additionally, these studies predict that Hit 2 binds less efficiently to CUG repeat RNAs than to CAG repeat RNAs.

### Lead candidate compound Hit 2 reduces expression of CAG expansion transcripts across multiple patient derived fibroblast lines and the *Atxn1^154Q/2Q^*SCA1 mouse model

To investigate the potential of Hit 2 as a regulator of CAG expansion transcripts in the context of SCA disease causing genes, we treated SCA1, SCA3 and SCA7 patient derived fibroblast lines with Hit 2 for 48 hours. Again, we initially set out to identify a dose range that did not negatively impact cell viability. In three control lines, one SCA1, two SCA3 and one SCA7 fibroblast line, Hit 2 was well tolerated up to 10µM. At 25µM, there was an overall trend of reduced cell viability with Hit 2 significantly reducing viability by 27.1% for Control 2 (*P*=0.0047) compared to DMSO treatment (Fig 5A). In the SCA1 patient derived cell line, 100nM Hit 2 reduced expression of *ATXN1* by 73.2% (*P*=0.0013) and at 1µM Hit 2, *ATXN1* expression was reduced by 59.9% (*P*=0.0036; Fig 5B). For SCA3-1, a 19.3% reduction in *ATXN3* expression was seen at 10nM (*P*=0.0186; Fig 5C) and for SCA3-2, at 1nM, 10nM and 100nM Hit 2 reduced *ATXN3* expression by 24.2% (*P*=0.0246), 32.7% (*P*=0.0046) and 33.9% (*P*=0.0037), respectively (Fig 5D). In two of the three control fibroblast lines, no effect on *ATXN1* expression with Hit 2 was observed. For the third cell line (Control 3) a 12.8% reduction of *ATXN1* expression was observed at 100nM (*P*=0.0424; Fig S4A). No measurable reduction of *ATXN3* expression was observed at 10nM or 100nM concentration in the three control cell lines (Fig S4B). Together, these data indicate that Hit 2 regulates CAG transcript levels in the context of multiple SCA disease genes and does so in a repeat expansion selective manner.

**Figure 5.**
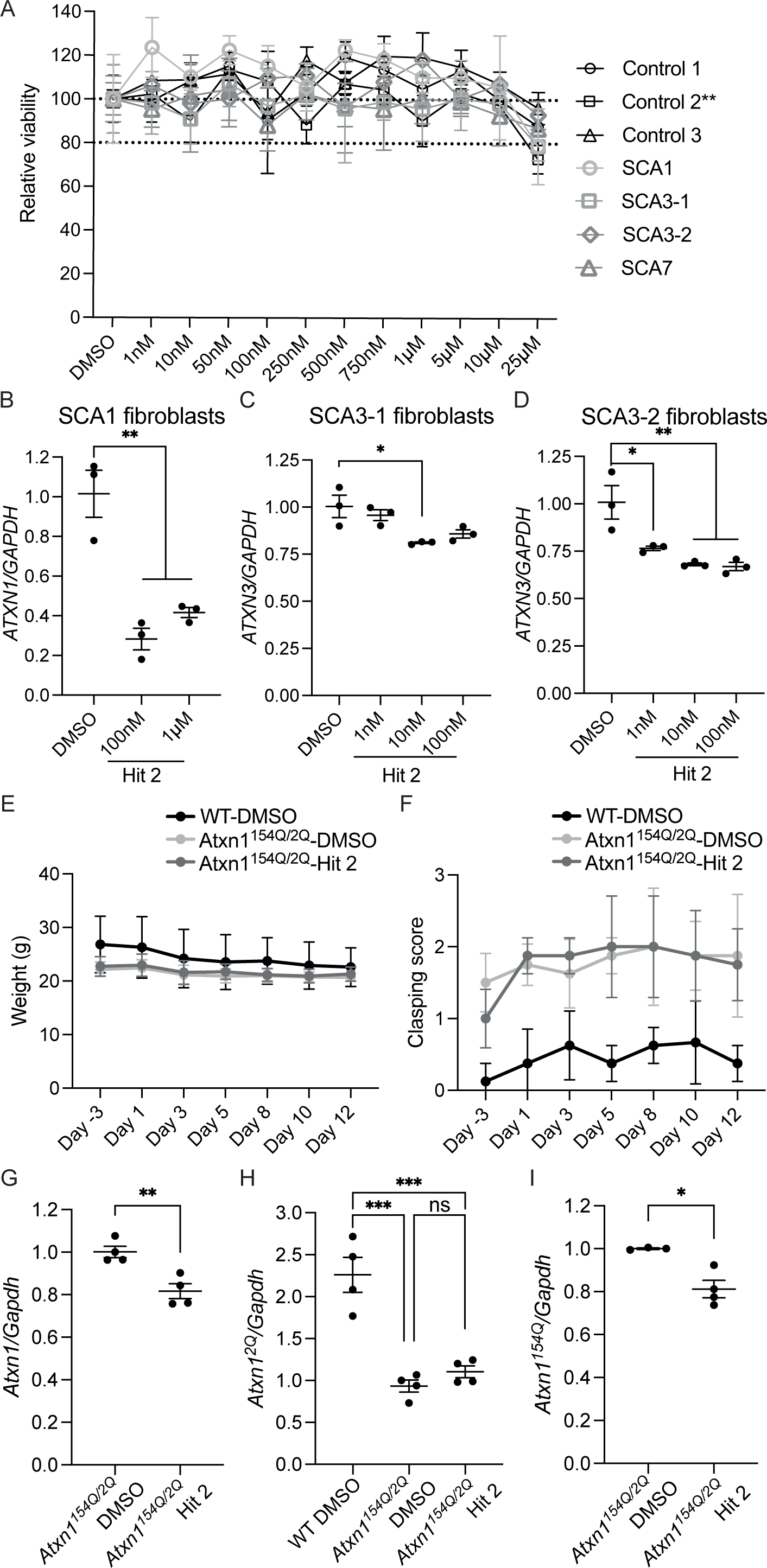
Hit 2 reduces expression of CAG expansion transcripts in SCA1 and SCA3 patient derived fibroblast lines and in an SCA1 mouse model. **A** Relative cell viability of three control, one SCA1, two SCA3 and one SCA7 fibroblast lines following 48 hours treatment with Hit 2; dotted lines indicate 100% and 80% relative viability; n=4, mean±SD, two-way ANOVA with Dunnett’s multiple comparisons test; Control 2 shows a significant reduction in viability at 25µM compared to DMSO: ** *P*<0.01; all other cell lines show no significant difference between DMSO and 25µM. **B-D** Hit 2 reduces expression of *ATXN1* transcripts in SCA1 patient-derived fibroblasts (**B**), and *ATXN3* in two SCA3 fibroblast lines (**C, D**); n=3, mean±SEM; one-way ANOVA with Tukey’s multiple comparisons test, * *P*<0.05, ** *P*<0.01. **E, F** Weight (g) (**E**) and clasping score (scored between 0 [no phenotype] and 3 [full clasp]) (**F**) across the two week treatment regimen with baseline three days before initial injection; n=4, mean±SD, clasping score is an average of two independent investigators per mouse per timepoint. **G-I** Hit 2 reduces expression of *Atxn1* (**G**), specifically *Atxn1^154Q^* alleles (**I**) but not *Atxn1^2Q^* alleles in the *Atxn1^154Q/2Q^* SCA1 mouse model following a two week IP injection regimen from 18 to 20 weeks of age; n=4, mean±SEM; **G,I** unpaired two-tailed t-test, **H** one-way ANOVA with Tukey’s multiple comparison test, ns – not significant, * *P*<0.05, ** *P*<0.01, *** *P*<0.001.

As Hit 2 is predicted to bind CUG repeat RNA, although with lower affinity than CAG repeat RNA, we wanted to understand whether Hit 2 could reduce expression of CUG expansion transcripts in cell culture. To do this, we differentiated DM1 patient derived myoblasts into myotubes and performed a three-day treatment of Hit 2. Increasing doses of Hit 2 showed a trend to increase expression of *DMPK*, the CUG repeat expansion containing gene causative for DM1^1^, with 1µM Hit 2 increasing *DMPK* expression by 51.6% (*P*=0.0266; Fig S4C). To understand if Hit 2 is capable of rescuing CUG induced mis-splicing which is a transcriptomic hallmark of DM1^1^, we performed RT-PCR analysis and found that Hit 2 did not affect the splicing of three alternatively spliced events in DM1 myotubes (Fig S4D-F). This therefore demonstrates that the mechanism of action of Hit 2 is specific to CAG repeats; possibly indicating that Hit 2 does not bind CUG repeats strong enough to reduce expression of CUG repeat containing transcripts or to displace the MBNL RNA binding proteins responsible for CUG induced mis-splicing.

Given that Hit 2 was well tolerated in patient derived fibroblast cell lines and selectively reduced *ATXN1* and *ATXN3* expression levels in a repeat expansion selective manner, we investigated if Hit 2 was capable of reducing expression of CAG SCA disease associated genes *in vivo*. Because of the large effect size seen in SCA1 fibroblasts, we decided to perform the *in vivo* treatment of Hit 2 in an SCA1 mouse model and selected the *Atxn1^154Q/2Q^*knock-in model to ensure we were investigating *Atxn1* expression under endogenous control. To generate sufficient quantity of Hit 2 for an *in vivo* study, we synthesized Hit 2 in house as previously described^32^ (Fig S5). Animal model studies using Hit 2 have not previously been reported and so with this study, we set out to perform a short two-week treatment regimen at 25mg/kg (<4ml/kg DMSO) and to closely monitor the mice for signs of toxicity during this time frame. Starting at 18 weeks of age, Hit 2 or DMSO was injected via IP for 5 days, followed by two rest days and then an additional 5 days of IP injection before sacrificing the mice. Across the treatment timeline, Hit 2 did not negatively impact mouse weight beyond the effects seen for DMSO alone (Fig 5E) and there was no negative impact of Hit 2 on the clasping phenotype of *Atxn1^154Q/2Q^*mice compared to DMSO injected mice (Fig 5F). Together these data provide the first indication that Hit 2 is well tolerated *in vivo* over a short study time frame at 25mg/kg.

Following completion of the two-week injection regimen, we performed RNA extraction on cerebellum from Hit 2 and DMSO treated SCA1 and WT mice. Compared to DMSO injected *Atxn1^154Q/2Q^*mice, Hit 2 reduced expression of *Atxn1* in the cerebellum by 18.5% (*P*=0.0056; Fig 5G). To understand if this reduction was specific for the expansion allele, we utilized allele specific qPCR primers to differentiate between the 2Q and 154Q alleles. As expected, DMSO injected WT mice showed approximately twice the expression of *Atxn1^2Q^* compared to SCA1 mice: 132.6% increase compared to *Atxn1^154Q/2Q^*-DMSO (*P*=0.0002) and 115.7% increase in *Atxn1^2Q^* compared to Hit 2 treated SCA1 mice (*P*=0.0005), consistent with previous studies^33^. There was however no difference in *Atxn1^2Q^* expression between DMSO and Hit 2 injected SCA1 mice (Fig 5H). This is in contrast to the *Atxn1^154Q^*allele selective qPCR which showed a 18.8% reduction in expression of *Atxn1^154Q^* in *Atxn1^154Q/2Q^*-Hit 2 mice compared to DMSO injected (*P*=0.0109; Fig 5I). Together these data demonstrate that over a two-week period, Hit 2 is well tolerated and can selectively reduce expression of expansion *Atxn1* alleles *in vivo*.

### Hit 2 rescues dysregulation of alternative splicing in the *Atxn1^154Q/2Q^* SCA1 mouse model

To understand more about the effects of Hit 2 *in vivo* we performed RNA-Seq on cerebellum from WT-DMSO, *Atxn1^154Q/2Q^*-DMSO and *Atxn1^154Q/2Q^*-Hit 2 treated mice (Table S1). Recently, we^33^ and others^34^ reported that dysregulation of alternative splicing is a widespread transcriptomic hallmark of CAG expansion SCAs including in the *Atxn1^154Q/2Q^*mouse model. Furthermore, we demonstrated that alternative splicing could be used as a target engagement biomarker for CAG SCAs^33^. We therefore set out to assess the effect of Hit 2 on both gene expression and alternative splicing. Because previous alternative splicing analyses in CAG SCAs were performed on datasets generated using polyA selection^33,34^ and our standard pipeline for alternative splicing analysis is to use rRNA depletion, we first assessed whether our in house dataset recapitulated the published alternative splicing studies. Of the significant mis-spliced events between WT-DMSO and *Atxn1^154Q/2Q^*-DMSO mice with a change in percent spliced in (ΔPSI) >10% and FDR<0.1, skipped exon (SE) events were the most frequently dysregulated, accounting for 50.1% of all events (Fig S6A). Dysregulation of both inclusion and exclusion events were detected with a mean ΔPSI for inclusion events of 23.0% and a maximum ΔPSI of 92.0%; for exclusion events these values were 19.4% and 58.4% respectively (Fig S6B). Gene ontology enrichment analysis revealed that skipped exon events occurred in genes implicated in nuclear processes (eg GO:0051647), synapse organization (eg GO:0050807), calcium ion regulation and membrane potential (eg GO:0048791 and R-MMU-112308; Fig S6C). While all these findings are consistent with the extent of splicing dysregulation and the pathways affected previously reported^33,34^, we also confirmed dysregulation of *Trpc3* exon 9 (Fig S6D) and *Anks1b* exon 5 (Fig S6E) SE events that were previously reported^33,34^. Together, these data independently validate that dysregulation of alternative splicing is a prominent transcriptomic feature affecting disease relevant pathways in CAG expansion SCAs.

We next sought to understand whether Hit 2 impacted alternative splicing and differential gene expression in the *Atxn1^154Q/2Q^*mouse model. Because SE events accounted for 50.1% of all miss-spliced events, we focused our alternative splicing analysis on SE events. Of the 495 SE events dysregulated between WT-DMSO and *Atxn1^154Q/2Q^*-DMSO mice, the majority showed smaller differences between WT-DMSO and *Atxn1^154Q/2Q^*-Hit 2 with a trend towards overall rescue of events. We also detected 273 SE events dysregulated between WT-DMSO and *Atxn1^154Q/2Q^*-Hit 2, but not between WT-DMSO and *Atxn1^154Q/2Q^*-DMSO; demonstrating an off-target effect of Hit 2 accounting for 0.59% of all SE events detected (Fig 6A). Interestingly, far fewer differentially expressed genes were detected (20 genes; log_2_FC>1, padj<0.05) with most events clustering around the dotted line indicating no rescue. Four events were differentially expressed between WT-DMSO and *Atxn1^154Q/2Q^*-Hit 2 but not WT-DMSO and *Atxn1^154Q/2Q^*-DMSO indicating minimal off target effects of Hit 2 on differential gene expression (Fig 6B). This limited rescue of differential gene expression compared to strong rescue of alternative splicing by Hit 2 is exemplified by PCA plots based on significant events between WT-DMSO and *Atxn1^154Q/2Q^*-DMSO (Fig 6C, D). For differential gene expression, there is clear separation between WT and SCA1 mice but overlap between the Hit 2 and DMSO injected SCA1 mice (Fig 6C) indicating a lack of rescue of differentially expressed genes which is also clearly seen at the level of individual events (Fig 6E). In contrast, the PCA plot for skipped exons shows clear separation between all three treatment groups along the x-axis accounting for 36.9% of variance, with *Atxn1^154Q/2Q^*-Hit 2 mice approximately half-way between WT-DMSO and *Atxn1^154Q/2Q^*-DMSO mice (Fig 6D). These data therefore show that Hit 2 rescues dysregulation of alternative splicing in *Atxn1^154Q/2Q^*mice with limited off target effects and without impacting differential gene expression.

**Figure 6.**
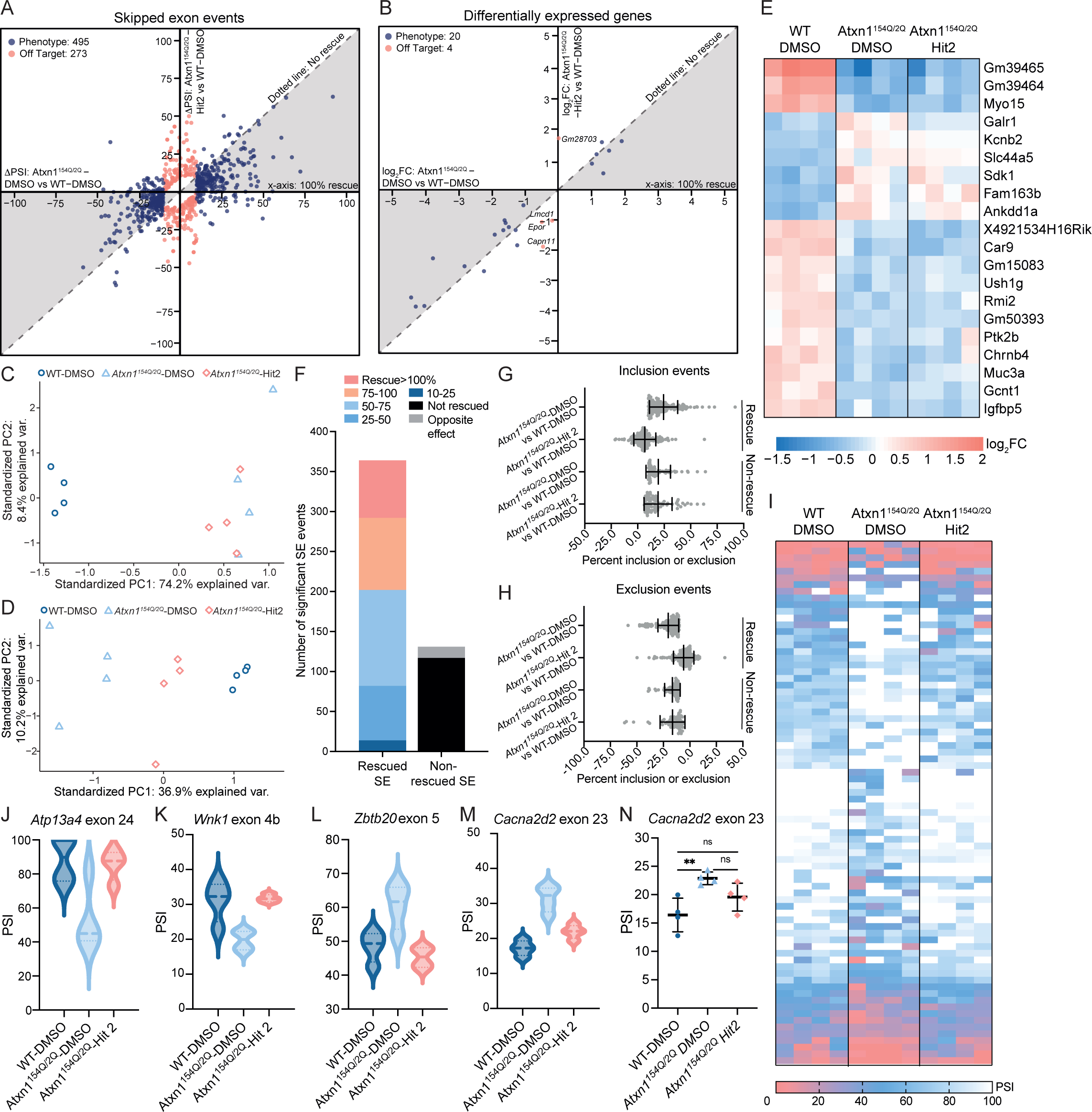
Lead candidate Hit 2 rescues alternative splicing dysregulation in the *Atxn1^154Q/2Q^* SCA1 knockin mouse model. **A, B** ΔPSI (**A**) and log_2_FC (**B**) for WT-DMSO vs *Atxn1^154Q/2Q^*-DMSO (*x*-axis) and WT-DMSO vs *Atxn1^154Q/2Q^*-Hit 2 (*y*-axis) showing events significantly different between WT-DMSO vs *Atxn1^154Q/2Q^*-DMSO (ΔPSI>10%, FDR<0.1; log_2_FC>1, padj<0.05; Phenotype – Blue) and WT-DMSO vs *Atxn1^154Q/2Q^*-Hit 2 (Off Target – Pink). Events that cluster around the *x*-axis show 100% rescue on Hit 2 treatment and events that cluster along the dotted line show no-rescue no Hit 2 treatment. **C, D** PCA plots based on phenotype events (see Fig 6A and B) for differential gene expression (**C**) and alternative splicing (**D**) showing clear rescue of dysregulation of alternative splicing but not differential gene expression on Hit 2 treatment. **E** Heatmap showing 20 genes differentially expressed between WT-DMSO and *Atxn1^154Q/2Q^*-DMSO (log_2_FC>1, padj<0.05) and their response to Hit 2 treatment. **F** Number of SE events rescued (>10% rescue and ΔPSI>5% between SCA DMSO and Hit2) and number of non-rescued SE events which include events showing an opposite effect (>10% change in splicing away from WT PSI with a minimum ΔPSI of 5% between SCA DMSO and Hit2) and events that are not-rescued (all remaining events). **G, H** Percentage of exon inclusion (positive) or exclusion (negative) for events that show increased exon inclusion between WT-DMSO and *Atxn1^154Q/2Q^*-DMSO (**G**) and events that show increased exon exclusion between WT-DMSO and *Atxn1^154Q/2Q^*-DMSO (**H**) (both: FDR<0.1, ΔPSI>10%,) and the response to Hit 2 treatment, separated into rescued and non-rescued events (based on Fig 6F), mean ± SD. **I** Heatmap showing SE events dysregulated between WT-DMSO and *Atxn1^154Q/2Q^*-DMSO (ΔPSI>10%, FDR<0.1) and their response to Hit 2 treatment for events significantly rescued by Hit 2 (*Atxn1^154Q/2Q^* –DMSO vs –Hit 2: ΔPSI>5%, FDR<0.1, >10% rescue). **J, K, L, M** Violin plots of RNA-Seq data for SE events significantly dysregulated between WT-DMSO and *Atxn1^154Q/2Q^*-DMSO and significantly rescued following Hit 2 treatment, showing that the increased exclusion of *Atp13a4* exon 24 (**J**) and *Wnk1* exon 4b (**K**) and the increased inclusion of *Zbtb20* exon 5 (**L**) and *Cacna2d2* exon 23 (**M**) is rescued on Hit 2 treatment, line indicates median. **N** RT-PCR analysis of *Cacna2d2* exon 23 showing rescue following Hit 2 treatment; n=4, one-way ANOVA with Tukey’s multiple comparisons test, ns – not significant, ** P<0.01, mean±SD.

To further understand the extent of alternative splicing rescue, we categorized the 495 SE events (Fig 6A, S6A) based on the size of change (percent rescue) between *Atxn1^154Q/2Q^* –DMSO and –Hit 2. Events were classified as rescued if the PSI for *Atxn1^154Q/2Q^*-Hit 2 changed >10% in the direction of the control mice and had a ΔPSI>5%, compared to *Atxn1^154Q/2Q^*-DMSO. Events were classified as changed in the opposite direction if ΔPSI>5% for *Atxn1^154Q/2Q^*-Hit 2 versus – DMSO and there was >10% shift in the opposite direction to the control mice; the remainder of events were classified as not rescued. Of the 495 SE events 364 (73.5%) were rescued with 282 rescued by >25%; only 14 events (2.83%) showed an opposite effect while 117 events (23.6%) remained unchanged (Fig 6F). For rescued inclusion events, the mean ΔPSI compared to WT-DMSO reduced from 24.4% for *Atxn1^154Q/2Q^*-DMSO to 6.73% for *Atxn1^154Q/2Q^*-Hit 2 with a corresponding ∼30% change in PSI for the minimum and maximum PSI values (Fig 6G). Likewise, for rescued exclusion SE events, the mean ΔPSI shifted from –20.4% for WT-DMSO vs *Atxn1^154Q/2Q^*-DMSO to –5.83% for WT-DMSO vs *Atxn1^154Q/2Q^*-Hit 2 with an 8.55% shift in the minimum ΔPSI and a 43% shift in the maximum ΔPSI (Fig 6H). Next, we further filtered these SE events to identify events significantly alternatively spliced between *Atxn1^154Q/2Q^* –DMSO and –Hit 2 mice, indicating significant rescue of these events (Fig 6I). Many of these events affect genes involved in disease relevant pathways, for example cellular calcium ion homeostasis (*Atp13a4*, Fig 6J), a serine/threonine protein kinase with links to hereditary neuropathy (*Wnk1*, Fig 6K), transcription factors (*Zbtb20*, Fig 6L), and a voltage-dependent calcium channel linked to cerebellar atrophy (*Cacna2d2*, Fig 6M). Using RT-PCR, we confirmed that *Cacna2d2* exon 23 shows significantly increased inclusion at 12 (5.96%, *P*=0.0005) and 23 (8.77%, *P*=0.0004) weeks of age in *Atxn1^154Q/2Q^* mice compared to WT mice (Fig S6F) and that this event is rescued following Hit 2 treatment (Fig 6N). In summary, Hit 2 reduces the severity of the transcriptomic phenotype in *Atxn1^154Q/2Q^* mice with minimal off target effects.

## Discussion

There is currently a large unmet therapeutic need for spinocerebellar ataxias caused by CAG repeat expansions. Despite recent progress in preclinical therapy development^3,4^, and ongoing clinical trials using ASOs for these devastating diseases^2^, there remains a gap in the development of small molecule therapeutics that can target the shared pathogenic source, the CAG repeat expansion, across this class of SCAs. This is reflected in the lack of small molecules published to reduce expression of CAG expansion transcripts in SCAs.

Here, to address this gap, we developed a disease-gene independent, mechanism-agnostic CAG repeat selective small molecule screening cell system that recapitulated phenotypic and transcriptomic hallmarks of mouse models of CAG expansion SCAs. Utilizing this cell line, we identified that the FDA approved microtubule inhibitor, colchicine was capable of selectively reducing expression of CAG expansion transcripts relative to a zero-repeat control. In SCA1, SCA3 and SCA7 patient derived fibroblasts, colchicine was able to reduce expression of the disease-causing transcripts in a repeat dependent manner. By screening 1584 structurally diverse small molecules, we identified a pyrazole based compound, Hit 2 (IUPAC name: 4-[(4-methylphenyl)diazenyl]-1-phenylpyrazole-3,5-diamine; CAS number: 5456-92-8), that selectively reduced expression of CAG expansion transcripts and polyglutamine expansion proteins relative to zero repeat controls. Hit 2 reduced levels of *ATXN1* and *ATXN3* transcripts in SCA1 and SCA3 patient derived fibroblast lines but not in control fibroblast lines. Hit 2 was also shown to selectively reduce expression of expansion alleles and rescue disease-associated alternative splicing dysregulation in a knock-in mouse model of SCA1 with minimal off target effects on the transcriptome. This first in mouse study of Hit 2 was therefore able to independently validate our recent data demonstrating the feasibility and validity of assessing alternative splicing dysregulation as a target engagement biomarker in CAG expansion SCAs^33^. Interestingly, colchicine and Hit 2 were identified to regulate CAG expansion transcripts through distinct mechanisms with Hit 2 predicted to bind to CAG helical RNA. This therefore demonstrates that our novel small molecule screening system is capable of identify compounds with different mechanisms of action for regulating expression of CAG repeat expansions.

Previous mechanism-agnostic small molecule screening approaches have also identified compounds with therapeutic promise in SCAs, primarily focusing on SCA3^35–38^. Multiple cell-based screening systems have been developed to identify compounds capable of regulating expression of ATXN3 protein based on a luciferase readout^35,37^. One of these screening systems identified three compounds, including the FDA approved atypical antipsychotic agent aripiprazole, capable of reducing expression of human ATXN3-84Q and mouse Atxn3 protein in organotypic brain slice cultures from 84Q SCA3 transgenic mice^35^. Interestingly however, another cell based screening system did not identify compounds capable of reducing ATXN3 expression but found that statins upregulate expression of *ATXN3* mRNA and protein levels; providing valuable insight into management of SCA3^37^. Other screens in *Caenorhabditis elegans* models of SCA3 identified compounds capable of improving behaviour phenotypes^36,38^ and found that the FDA approved selective serotonin reuptake inhibitor, citalopram was capable of reducing mutant ATXN3 aggregation, and expression in some brain regions, in SCA3 mouse models^38,39^. While these screening systems highlight the ability of mechanism agnostic screening approaches to identify compounds that can regulate polyglutamine containing protein behaviour, many of these compounds act on both normal and expanded ATXN3 and do not reduce expression of *ATXN3* mRNA.

In contrast to these studies, our screening approach set out with a broader aim to identify compounds with therapeutic potential across CAG SCAs. Other groups have sought to identify candidate therapeutics that could be used across multiple CAG SCAs through their ability to rescue cellular processes commonly disrupted in SCAs. For example, as Purkinje cell function is impacted in many SCAs, therapies that target this, such as 4-aminopyridine and chlorzoxazone^40–43^, may provide therapeutic benefit across a range of SCAs. Similarly, Trehalose, a compound that has been shown to induce autophagy, has shown promise in models of SCA3 and SCA17^44,45^, and is currently in clinical trials for SCA3^2^. Additionally, more general neuroprotective compounds such as riluzole and its prodrugs are in clinical trials for multiple CAG SCAs, but so far have shown mixed results^2^. While these approaches may result in therapeutic benefit across multiple SCAs, the window of benefit is likely restricted due to the ongoing neurodegenerative effects caused by the disease-causing genes which cannot be targeted using these approaches.

In this study, we were able to identify compounds that target the pathogenic source of CAG expansion diseases *and* may yield therapeutic benefit across multiple CAG SCAs. By integrating a repeat-selective aspect into a gene-context independent screening system we selected for compounds that retained repeat selectivity in more complex and disease relevant model systems of multiple CAG SCAs. Despite these strengths, our novel screening system has limitations: primarily a lack of ability to control expression from the integrated constructs. In combination with the high level of cell toxicity associated with expressing CAG repeats in this system, the lack of ability to control expression results in selective pressure against both the length and the expression level of the pure CAG repeat. Reengineering this system into an inducible system would reduce the selective pressure against expression of the integrands and increase the longevity of an individual clonal cell line as well as providing a more tractable model system for investigating the effects of expression of CAG repeat expansions.

Multiple previous screening systems in CAG SCAs^35,38,39^, and indeed our own study, identified FDA approved compounds with therapeutic potential in CAG SCAs. Repurposing of FDA approved compounds is designed to accelerate the pace of therapeutic development by advancing candidates ready for human trials that do not require extensive pharmacodynamic and/or toxicity studies. Here we identified the FDA approved microtubule regulating compound colchicine as a CAG repeat containing transcript regulating compound. One of the draw backs of colchicine in this study is that it was unable to reduce expression of the polyQ proteins in our screening cell line. This raises the idea of combinatorial therapies based on targeting different levels of the pathogenic cascade or combining therapies with different mechanisms of action to reduce the dose of each therapy and thereby reduce unwanted toxicity. In the context of combining small molecule approaches with ASOs this could potentially help overcome issues of toxicity^2,14^ whilst retaining effect size of reducing CAG expansion containing transcripts. If paired with non-allele selective ASO approaches, repeat selective small molecules could also aid in preventing excessive reduction of expression of the non-expanded protein levels. Similarly, given that both the FDA approved compounds citalopram and aripirazole have been shown to reduce levels of ATXN3 protein but not mRNA^35,38,39^, it would be interesting to understand if, at least in the context of SCA3, there was an additive effect of either compound in combination with colchicine, or Hit 2, which regulate CAG containing mRNA.

Interestingly, another microtubule regulating compound, nocodazole, was identified through a screen in SCA6 for its ability to prevent internal ribosome entry site (IRES) mediated translation of the α1ACT protein product which is toxic in the context of SCA6^46^. It remains to be seen whether nocodazole also regulates CAG expansion expression in a similar manner to colchicine. However, the rationale behind using microtubule regulating compounds in neurodegenerative diseases in which the cytoskeleton and microtubule associated processes are disrupted is questionable^33,47,48^ and may accelerate degeneration through distinct mechanisms rather than slow the disease process. Furthermore, although other microtubule regulating compounds may be tolerated better, colchicine is typically prescribed on a short-term basis due to the effects of colchicine poisoning^49^. This is a toxicity syndrome that can lead to death and in some cases causes neuropathy, myopathy, distal sensory abnormalities, and nerve conduction impairment^49^ and so colchicine does not represent a good therapeutic for neurodegenerative conditions.

Hit 2 represents a promising starting point for developing a novel therapeutic compound for CAG repeat expansion diseases because it is capable of reducing both CAG expansion transcript and polyQ protein expression. Despite the current problems of insolubility of Hit 2 and the requirement of DMSO or methanol as a solvent, understanding the binding mode and mechanism of action of Hit 2 will enable us to perform structure activity relationship studies to identify structural analogs of Hit 2 with improved solubility and more druglike properties. Although our first in mouse experiment of Hit 2 demonstrated that it was well tolerated over a two-week period, the mice in the study did not gain weight due to the daily DMSO administration, and so a compound with improved solubility will be required before a long-term behavior and survival study can be performed. Despite this, our short-term study demonstrated promising effects of reducing *Atxn1* expression levels in a repeat selective manner and rescuing transcriptomic hallmarks of disease. During development of Hit 2, it will be important to consider the effects on CUG expansion transcripts due to the known role of antisense transcripts in repeat expansion diseases^50–52^. Here we found that Hit 2 increased expression of *DMPK* in DM1 myotubes. If this increase in expression also occurs for antisense CUG expansion transcripts in CAG SCA disease causing genes, this could lead to disease perpetuating off-target effects. Indeed, future studies will be needed to understand whether antisense CUG expansion containing transcripts are expressed in the *Atxn1^154Q/2Q^* mouse model and whether some of the off-target effects of Hit 2 on alternative splicing seen are due to upregulation of these antisense transcripts.

Overall, this study provides proof of concept that our disease-gene independent, mechanism agnostic small molecule repeat selective screening system is able to identify compounds that regulate CAG expansion transcript abundance in a repeat length dependent manner across multiple CAG expansion diseases. Through this study we have reported the first compounds capable of reducing expression of CAG expansion containing transcripts including our promising lead candidate, Hit 2, which warrants further investigation to improve its solubility and druglike properties. This work also paves the way for further screening studies in a similar repeat selective screening system to identify compounds with the potential for accelerated pace of therapeutic development such as FDA approved compounds. Together this study lays a strong foundation for the potential for repeat selective small molecules as shared therapeutics, either alone or in combination with other therapeutic approaches, across multiple CAG repeat expansion spinocerebellar ataxias, and potentially across the broader class of CAG microsatellite expansion diseases.

## Acknowledgements

The authors thank the RNA Institute Summer Bioinformatics program for funding and training undergraduate trainees who contributed to this work and members of the Berglund, Reddy and Shin laboratories for their valuable input and discussion. The authors would like to thank Renjie Song, Director of the Immunology Core at the New York State Department of Health Wadsworth Center for performing single cell sorting to generate the clonal screening cell lines. This work was supported by funding from the National Ataxia Foundation, NIH K99 NS124994 (H.K.S.), RNA Institute Summer Bioinformatics Fellowships (V.A.D., C.C.D., A.K.M.), NIH R21 NS114794 (D.S.S.), NIH P50 NS04843 (J.A.B.), NIH R01 NS120485 (J.A.B., J.D.C. and K.R.), and NIH R01 NS135254 (J.A.B., H.K.S. and D.S.S.).

## Author contributions

HKS and JAB conceived the study and wrote the paper. HKS and JAB designed experiments and interpreted the data. HKS, JAB, JDC and KR designed the small molecule screening system, screening approach and interpreted the data. HKS generated the small molecule screening cell line and performed the Diversity Set VI screen and validation. HKS, JAB, DSS and JDC designed the Hit 2 treatment experiment of *Atxn1^154Q/2Q^* mice and interpreted data. DSS maintained *Atxn1^154Q/2Q^*mouse model, HKS performed the Hit 2 treatment study, VAD and CDL analysed clasping behaviour. AKM characterised the cell toxicity phenotype and confirmed CAG repeat length in clone 10, 15 and 37. HKS and JAF identified colchicine as a CAG repeat regulating compound. AA performed SCA patient derived fibroblast experiments, HKS performed control fibroblast experiments, JAF performed DM1 and control myotube studies.

HM synthesized Hit 2 and performed mass spectrometry. SV performed molecular docking studies. AA and SS generated RNA-Seq libraries and VAD performed RNA-seq analyses and validation for GSE249555. AA generated RNA-Seq libraries, CDL, CCD and HKS performed RNA-seq analyses and AA performed validation for GSE249556. HKS performed statistical analyses. All authors reviewed the manuscript and provided input.

## STAR Methods

### Data and material availability

All datasets used in this study are publicly available through the NCBI Gene Expression Omnibus (GEO) under the BioProject accession number PRJNA1049475 using the following GSE numbers: GSE249555 and GSE249556 (Table S1). In house synthesized Hit 2 and the plasmids and protocols used to generate the stably transfected ATG-(CAG)_60_-Myc-NLuc/ATG-(CAG)_0_-Myc-FLuc HEK293T cell line are available on request. Further information and requests for resources and reagents should be directed to and will be fulfilled by the lead contact, J Andrew Berglund.

### Generation and culture of stable CAG repeat expressing HEK293T screening cell line

Stable HEK293T cell lines expressing ATG-(CAG)_60_-Myc-NLuc and ATG-(CAG)_0_-Myc-FLuc were generated as previously described^21^. Gene blocks containing the ATG-(CAG)_60_-Myc-NLuc-Pest and ATG-(CAG)_0_-Myc-FLuc-Pest cassettes were ordered from Genscript. These were cloned into the constructs we previously used to generate our DM1 screening system^21^ between the EF1α promoter and bGH poly(A) signal using the MluI and KpnI sites such that the *DMPK* containing cassette was removed and the PiggyBac transposon terminal repeats, puromycin selection cassette, EF1α promoter and bGH poly(A) signal were retained. The MluI site was blunted and thus destroyed during the cloning process. These constructs were transfected into HEK293T cells (ATTC) using lipfectamine 3000 (Invitrogen) together with the PiggyBac mPB transposase and selected by puromycin (Gibco). Cells were suspended in a solution of 2% dialyzed HI-FBS (Cytiva), 10mM HEPES (Sigma) in 1xPBS and single cells were isolated by flow cytometry using a BD FACSAria IIu and colonies were cultured in 96-well plates. Colonies were expanded and subjected to luciferase assay and multiplex qPCR to identify clonal lines that express both ATG-(CAG)_60_-Myc-NLuc and ATG-(CAG)_0_-Myc-FLuc. Parental and clonal cell lines were cultured in DMEM (Corning) supplemented with 10% heat-inactivated fetal bovine serum (HI-FBS; Corning) and 1x penicillin/streptomycin (ThermoFisher) in a humidified atmosphere at 37°C at 5% CO_2_; whilst cells expanded from single cells into clones, they were cultured in the same media but with 20% HI-FBS.

### Transient transfections of siRNAs and DNA plasmids

(CAG)_7_ siRNA and scrambled control siRNA (Dharmacon) were transfected for a final concentration of 17.5nM using INTERFERin® (Polyplus) in Opti-MEM (Gibco). For plasmid co-transfections, 100ng of each plasmid was transfected into parental HEK293T cells in a 12 well plate using Lipofectamine 3000 (Invitrogen) in Opti-MEM (Gibco).

### Diversity Set VI small molecule library screen

Approximately 2.5 × 10^4^ cells per well were plated in a 96 well plate and cultured overnight in 100µL of DMEM supplemented with 10% HI-FBS and 1xP/S under standard conditions of 37 °C and 5% CO_2_. The next day, medium was removed and replaced with fresh medium (100µL) containing small molecule at a final concentration of 1µM or 100nM or DMSO as a control. All treatments occurred at 0.01% DMSO. The Diversity Set VI library from NCI DTP repository was screening in its entirety in clone 37 with subsequent validation screens of initial hits in clones 15 and 37. Each screening plate also contained two wells of each control: DMSO only, 0.1nM Colchicine, scrambled siRNA and (CAG)_7_ siRNA. Following 24 hours of treatment, media was removed, and cells were stored at −80 °C. The following day, plates were thawed, and each well was incubated with 20µL of 0.25% Igepal in buffer containing 10 mM Tris·HCl (pH 7.5) and 150 mM NaCl on an orbital shaker for 10 mins at room temperature; 4µL of “lysate” was then utilized for cDNA generation.

### Cell viability analysis and luciferase assay

Cell viability was quantified using the MultiTox-Glo Multiplex Cytotoxicity Assay (Promega) following the manufacturers’ instructions. For effects of small molecule treatments on cell viability, data is reported from the live cell component of the assay normalised to DMSO only for each cell line. For assessing cell toxicity of clone 10, 15 and 37, data from the dead cell assay component was normalised to live cells per well, and then to the average for parentals, to account for the difference in growth rate of the cell lines. PolyQ-Nluc and zero repeat Fluc were quantified using the Nano-Glo® Dual-Luciferase® Reporter Assay System (Promega), following manufacturers’ instructions. The Nluc reading was normalised to Fluc per well and then normalised to the average of DMSO only or scrambled siRNA negative control.

### Cell culture and treatment of CAG SCA patient derived fibroblast lines

All control and patient derived fibroblast cell lines (see key resources table) were cultured in Eagle’s Minimum Essential Medium (EMEM; Corning) containing 15% HI-FBS and 1xP/S at 37 C and 5% CO2. The cells were expanded and seeded in 12-well tissue culture plates with a density of approximately 3×10^4^ cells/ml. Once the cells reached ∼70% confluency, media was removed and replaced with fresh EMEM+FBS+P/S containing Hit 2 or colchicine at the specified concentrations or with a matched percentage of DMSO (vehicle; always ≤0.01% DMSO). Prior to adding fresh media containing DMSO or small molecules to the cells, the media/small molecule mix was vortexed for >20s. Following 48 hours treatment, media was removed, and RNA was extracted using Aurum mini kit (BioRad) with on-column DNase I treatment, following the manufacturer’s protocol.

### Culture and treatment of DM1 patient derived myotubes

To assess Hit 2 effects in a CTG repeat expansion disease, control and DM1 myoblasts were seeded in 12-well tissue culture plates in SkGMTM-2 BulletKitTM growth medium (Lonza) with a density of approximately 1×105cells/mL. Once myoblasts reached ∼80% confluency they were differentiated into myotubes for four days in DMEM/F12 50/50 (Corning) supplemented with 2% horse serum. Myotubes were treated with Hit 2 at the specified concentrations or with DMSO (vehicle) in the SkGMTM-2 BulletKitTM growth medium for three days. Cells were harvested after Hit 2 treatment and RNA was extracted using Zymogen’s Quick-RNA Miniprep kit with on-column DNase I treatment following manufacturer instructions.

### Mouse studies

Mice used in this study were housed and treated in accordance with the NIH Guide for the Care and Use of Laboratory Animals and complied with the Albany Medical College Institutional Animal Care and Use Committee (IACUC) guidelines under approved animal care and use protocol numbers 20-04002 and 23-03002. *Atxn1^154Q/2Q^* mice were originally obtained from Jackson Laboratories (strain number 005601) and maintained according to established breeding protocols with genotyping performed by PCR, at weaning and retroactively, following existing protocols^53^.

Starting at 18 weeks of age, mice were treated with 25mg/ml Hit 2 dissolved in DMSO or DMSO alone (DMSO <4ml/kg) via intraperitoneal (IP) injection daily for 5 days, followed by two rest days and an additional 5 days of IP injection. All injections were performed between 8 and 9am in the morning and on the final day of injections (20 weeks of age), after 3pm, mice were anesthetized using urethane in saline (1.2-1.5 g/kg) followed by a double thoracotomy and perfusion through the ascending aorta with 20 mLs 1x PBS. Cerebellum was removed and stored at –70°C. Three days prior to the first injection day and then on days 1, 3, 5, 8, 10 and 12, prior to IP injection, mice were weighed and suspended via their tail for a 10s video to allow for clasping phenotype to be assessed. The clasping phenotype was scored between 0 (no phenotype) and 3 (full clasp) by two independent investigators and the results averaged. The investigators who scored the clasping phenotype were not involved in the treatment regimen and were blinded to the day, genotype and treatment group of the mice. RNA from 6, 12, and 23 week WT and *Atxn1^154Q/2Q^*mice harvested as part of a previous study^33^ was utilized for validation of dysregulation of specific alternative splicing events in untreated mice across disease onset.

### Protein blotting

Media was removed from parental and clonal HEK293T cells which were then washed with 1xPBS and lysed in radioimmunoprecipitation assay (RIPA) buffer (Thermo Scientific) with 1X cOmplete Protease Inhibitors (Roche) for 15 min on ice. DNA was sheared by passage through a 21-gauge needle, lysates were centrifuged at 21,000 g for 15 min at 4°C, and the supernatant was collected. The protein lysate concentration was quantified using Pierce BCA Protein Assay Kit (Thermo Scientific), and 10μg of soluble protein lysates was separated on a 4–12% Bis–Tris gel (Bio-Rad) and transferred to a nitrocellulose membrane which was then blocked for 2 hours at room temperature in 5% dry milk in 1xPBS containing 0.05% Tween-20 (Sigma). The membrane was probed with primary antibodies (see Key Resources table) in blocking solution overnight at 4°C, washed three times in 1xPBS and incubated with species specific secondary antibodies (see Key Resources table; GAPDH was labelled with IRDye 680, all other antibodies were labelled with HRP conjugated secondary antibodies) in blocking solution for 1 hour at room temperature. Following three washes in 1x PBS at room temperature ProSignal Pico Spray (Prometheus) was used following manufacturers’ instructions and membranes were imaged using a ChemiDoc MP Imaging System (Bio-Rad).

### Reverse transcription

RNA concentrations were measured using nanodrop and 500ng total RNA was reverse transcribed using SuperScript IV reverse transcriptase (Invitrogen) with random hexamers (IDT) for patient derived fibroblast lines and qPCRs in Fig 1I, or RT-primer (see key resources table) for multiplex qPCR for CAG repeat containing cell lines. Reverse transcriptase reactions were performed with 200ng of input RNA for cDNA production for Control and DM1 myotubes.

### Quantitative PCR (RT-qPCR) analysis

Multiplex qPCR was carried out using Hot Start Taq 2× Master Mix (NEB) with Multiplex qPCR Fw and Multiplex qPCR Rv primers and Multiplex qPCR Probe 1 and Multiplex qPCR Probe 2 fluorescent probes (all IDT; see Key Resources table). For all other qPCRs cDNA was subjected to qPCR for 39 cycles with PowerUp SYBR Master Mix (Applied Biosystems) or Ssoadvanced Master Mix (BioRad) according to manufacturer’s instructions. All qPCRs were performed in a CFX384 or CFX96 Real-Time System (Bio-Rad). All qPCR reactions were performed in technical triplicates or quadruplicates using 1-3 uL cDNA depending on expression of target gene (i.e. 3uL input for GABRA3 qPCR, 1uL input for GAPDH). Ct values were obtained via CFX maestro software (Bio-Rad) and RT-qPCR data were analysed using the 2^-ΔΔCt^ method^54^. The levels of r(CAG)_60_ from small molecule treatments were normalized to r(CAG)_0_ and presented as relative mRNA levels by comparing to DMSO control treatments. GAPDH was used as the housekeeping gene. To confirm specificity of primers, qPCR was performed on RT reactions in which the RT enzyme was replaced with H_2_O (RT-) under the same conditions. All primers are listed in the key resources table.

### RT-PCR splicing analysis

PCR for selected splicing events was performed using the Taq 2x master mix (NEB) with 2 uL cDNA under the following conditions: 95°C 30s – 32 cycles of 95°C 30s, primer specific Tm 30s, 68°C 30s – 68°C 5 min. Primer sequences are included in the key resources table. Annealing temperatures and product sizes (inclusion, exclusion) are as follows: *MBNL1* exon 5 (Ta 58°C; 308bp, 250bp); *MBNL2* exon 5 (Ta 58°C; 266bp, 211bp); *SYNE1* exon 137 (Ta 58°C; 146bp, 83bp), *Cacna2d2* exon 23 (Ta 51°C; 107 bp, 86 bp). PCR products were resolved through capillary electrophoresis in a 5300 Fragment Analyzer system using the DNF-905 kit for 1-500 bp fragments (Agilent Technologies), following the manufacturer’s protocol. The relative fluorescence values (RFU) for the inclusion and exclusion bands, obtained from the ProSize 4.00 software (Agilent technologies), were used to calculate percent spliced in (PSI) for each exon of interest using the following formula: (Inclusion band RFU)/(Inclusion band RFU + Exclusion band RFU)*100. For alternative splicing analysis for *Atxn1^154Q/2Q^* and WT mice PCR products were resolved in technical triplicates on the fragment analyser and the average PSI is reported for each sample.

### RNA sequencing

RNA for CAG containing reporter cell lines was extracted using Aurum total RNA mini kit (BioRad) with on-column DNase treatment, following the manufacturer’s protocol. RNA for wild-type and KI mice was extracted using TRIzol (Ambion, Life Technologies), following the manufacturers protocol and a DNA digestion was performed using the TURBO DNA-free Kit (Invitrogen). The Qubit RNA high sensitivity assay was used to obtain RNA concentrations (Thermo Fisher Scientific). RNA quality was checked via capillary electrophoresis on a 5300 fragment analyzer (Agilent Bioanalyzer) using the RNA Analysis DNF-471 RNA kit (Agilent).

The NEBNext Ultra II Directional RNA Library Prep Kit (Illumina) with NEBNext rRNA Depletion Kit (New England Biolabs) was used to prepare RNA-seq libraries, with a total of 500 ng input RNA from each sample. The manufacturer’s protocols were followed, with the following exceptions: 40× adaptor dilutions were used, and 13 cycles of library amplification were performed. The resulting libraries were pooled in equimolar amounts, quantified using the NEBNext Library Quant Kit for Illumina (New England Biolabs), quality checked via capillary electrophoresis on the 5300 fragment analyzer (Agilent Bioanalzyer) using the DNF-474 HS NGS kit (Agilent). Libraries with sufficient quantity for RNA Seq of mouse cerebellum were loaded into P2 flow cell 1000/2000 by Illumina and were sequenced using paired-end, 100 base pair sequencing on the Illumina NextSeq 2000 sequencer at the University at Albany RNA Institute. Libraries with sufficient quantity for RNA Seq of parental, clone 15 and clone 37 cell lines sequenced using paired-end, 75 base pair sequencing (parental and clone 37), or 100 base pair sequencing (clone 15) on the Illumina NextSeq 500 the University at Albany Center for Functional Genomics. All datasets had total read depths greater than 40 million reads (Table S1).

FASTQ file quality was assessed using FastQC (version 0.11.9). Both paired-end files for samples 409, 422, 425, and 426 were downsampled using Seqtk (version 1.4) to either ∼65 or 70 million reads per file to more closely match the read depth of the other samples in the dataset. FASTQ files were then aligned to the GRCm39/mm39 mouse reference genome or GRCh38/hg38 genome using STAR with the –-quantModeGeneCounts option added to count the reads per gene (version 2.7.10a)^55^. Differential gene expression was performed in RStudio (2022.12.0; R 4.2.2) using DESeq2 (version 3.16)^56^ and genes that passed a threshold of padj<0.05 and log2FC>|1.0| were considered significantly differentially expressed. Volcano plots and heatmaps were created in RStudio (R version 4.2.2) using the packages EnhancedVolcano and pheatmap respectively. Differential gene expression events were considered off-target if they were significantly different between WT-DMSO and *Atxn1^154Q/2Q^*-Hit 2 (log2FC>|1.0| and Padj<0.05) but showed no difference between WT-DMSO and *Atxn1^154Q/2Q^*-DMSO (log2FC<|0.5|) mice. Alternative splicing analysis was performed using rMATS (version 4.1.2)^57^ and events were considered significantly misspliced if the false-discovery rate (FDR)<0.1 and ΔPSI>|0.1|. All ΔPSI values are converted from a ratio to a percentage with the threshold adjusted accordingly: ΔPSI>|10%|. Dysregulated skipped exon events were considered off-target if they were significantly differentially regulated between WT-DMSO and *Atxn1^154Q/2Q^*-Hit 2 or *Atxn1^154Q/2Q^*-DMSO and *Atxn1^154Q/2Q^*-Hit 2 mice (ΔPSI>|0.1| and FDR<0.1) but showed no difference between WT-DMSO and *Atxn1^154Q/2Q^*-DMSO (ΔPSI<|0.1| and P>0.05). Exon numbers are referred to by previously published exon numbers for the same coordinates or based on counting the first exon of the relevant transcript of a gene as exon 1. Coordinates for reported exons are as follows: *Trpc3* exon 9 (chr3:36688523-36688607), *Anks1b* exon 5 (chr10:90750557-90750629), *Atp13a4* exon 24 (chr16:29234600-29234697), *Wnk1* exon 4b (chr6:119905083-119905110), *Zbtb20* exon 5 (chr16:43392096-43392154), and *Cacna2d2* exon 23 (chr9:107396377-107396398). Gene ontology enrichment analysis was performed using Metascape (version v3.5.20230101)^58^.

### Computational modelling of Hit 2 binding to CAG and CUG RNA

To perform molecular docking studies for Hit 2 bound to CAG and CUG RNA, suitable RNA structures were identified from the PBD database. The RNA used for CAG repeats is a mixed sequence structure with a CAG helical region and a UUCG loop (PBD ID: 7D12)^59^. For CUG RNA, a CUG duplex was used (PBD ID: 1ZEV)^60^. These structures were energy minimized and molecular docking was performed using Molecular Operating Environment (MOE) (https://www.chemcomp.com/Products.htm). First, structure preparation, involved correcting protonation states and topologies of both the RNA and Hit2. Next molecular docking was performed using the triangle matcher and london dG method for placement of Hit 2 and scoring of the complexes respectively, to find the top 50 binding modes. These were then refined using induced fit method to allow for some local flexibility and scored using GBVI/WSA dG method to obtain the top 10 binding modes. The interaction between Hit 2:RNA complex structures were analyzed in MOE and visualized in PyMOL (The PyMOL Molecular Graphics System, Version 2.3.5, Schrödinger, LLC).

### Preparation of Hit 2

Hit 2 (IUPAC name: 4-[(4-methylphenyl)diazenyl]-1-phenylpyrazole-3,5-diamine; CAS number: 5456-92-8) was synthesized as previously described^32^. Briefly p-toluidine (1g, 1eq) was suspended in 2.5mL of 6N HCl (1.25mL of water and 1.25mL of concentrated HCl) and cooled in ice/water bath. Sodium nitrite (1.3eq in 5mL of water) was then added portion wise while stirring. This mixture was then added to a cold solution of sodium acetate (1.1eq), malononitrile (1eq) in 10mL of 50% aqueous ethanol. The thick precipitate was then stirred vigorously for 2hrs and filtered to afford a bright yellow solid, 853mg of which was carried onto the next step. Phenylhydrazine (1eq) was then added in ethanol and heated to 60°C overnight. This was then cooled and filtered to give a light brown solid, ∼900mg, MH+ 293.1979.

Figure S5B confirms the synthesized 4-[(4-methylphenyl)diazenyl]-1-phenylpyrazole-3,5-diamine (Hit 2) matched that of Figure S5A obtained from the NCI DTP (NSC: 21683; CAS: 5456-92-8); each compound was analyzed by the Agilent 6530B LC/MS QTOF mass spectrometer with dual ESI source. The fragmentor and skimmer voltage were set to 175 V and 65 V, respectively, while the voltage amplitude and capillary voltage were set to 750 V and 4000 V, respectively. The mobile phase was a mixture of water with 0.1% formic acid and acetonitrile (ACN) also containing 0.1% formic acid at gradient conditions:

| Time (min) | Water w/0.1% Formic Acid (%) | ACN w/0.1% Formic Acid (%) | Flow (mL/min) |
| --- | --- | --- | --- |
| 0 | 95 | 5 | 1 |
| 4 | 5 | 95 | 1 |
| 4.05 | 5 | 95 | 1.5 |
| 5 | 5 | 95 | 1.5 |
| 5.05 | 95 | 5 | 1 |
| 6 | 95 | 5 | 1 |

### Data analysis

For qPCR and alternative splicing validation analyses, statistical analysis was performed using GraphPad Prism 9. Grubb’s test with an alpha of 0.001 was used to identify and remove outliers for *Atxn1^154Q^* qPCR. Data are represented as mean ± standard error of the mean (SEM), mean ± standard deviation (SD) or median with range as appropriate which is listed in the figure legends. Statistical analyses were performed using two-tailed Student’s unpaired t-test, or one-way ANOVA with Tukey’s multiple comparisons test as appropriate and are listed in figure legends. A critical value for significance of p <0.05 was used throughout the study.

## Supplemental figure titles and legends

**Figure S1. RNA Sequencing analysis of Clone 15 and Clone 37 compared to Parental HEK293T cells**

**A, B** Clone 15 (**A**) and clone 37 (**B**) show more downregulated genes that upregulated genes; log_2_FC>0.5, *P*adj<0.05, n=3.

**C** Clone 15 and 37 show differential expression of genes previously reported to be differentially expressed in CAG expansion SCA mouse models, log_2_FC>0.5, *P*adj<0.05, n=3.

**D** Cycle threshold [C(t)] for *GABRA3*, *ALDH1L2* and *SLC7A11* which show an almost complete ablation of gene expression following analysis using the 2^-ΔΔCt^ method; *GAPDH* shown as housekeeping gene; n=6, mean±SEM.

**E, F** RNA-Seq log_2_fold change (log_2_FC) (**E**) and qPCR validation (**F**) for *METTL15*, included as a negative control for qPCR validation as this gene does not show differential expression on RNA-Seq analysis. **E** dashed line indicates log_2_FC=0.5, n=3. **F** n=6, mean±SEM, one-way ANOVA with Tukey’s multiple comparisons test, ns – not significant.

**Figure S2. Colchicine treatment does not reduce expression of CAG SCA genes in control fibroblasts**

**A-C** Colchicine does not affect expression of *ATXN1* or *ATXN3* in three control fibroblast lines and increases expression of *ATXN7* in two out of three control fibroblast lines at doses which reduce expression of equivalent expansion alleles (see Fig 2D-G); n=3, mean±SEM, unpaired two-tailed t-test, ns – not significant, * *P*<0.05, *** *P*<0.001.

**D, E** Colchicine does not reduce levels of polyQ-NLuc relative to FLuc in clone 37 (**D**) or clone 15 (**E**) at doses that result in a selective reduction of CAG expansion transcript levels; n=3, mean±SEM, one-way ANOVA with Tukey’s multiple comparisons test, ns – not significant.

**Figure S3. Positive controls and RT-qPCR assay controls for the diversity set VI screen**

**A, C** 24 hour treatment of clone 37 (**A**) and clone 15 (**C**) with CAG targeting siRNA selectively reduces (CAG)_60_ expression (normalized to (CAG)_0_); n=2 per screening plate, mean±SEM, unpaired two-tailed t-test, **** *P*<0.0001.

**B, D** Cycle threshold [C(t)] values for multiplex qPCR of DMSO negative control samples for reverse transcriptase (RT) reaction with RT enzyme (DMSO) and without RT enzyme in the reaction (DMSO RT-); n=2 per screening plate, mean±SEM.

**E** 24 hour treatment of clone 15 with 0.1nM colchicine selectively reduces (CAG)_60_ expression (normalized to (CAG)_0_); n=12 (n=2 per screening plate), mean±SEM, unpaired two-tailed t-test, **** *P*<0.0001.

**F** Screen results for each individual compound treatment in clone 15 on (CAG)_60_ normalized to (CAG)_0_, for the 72 hits identified in Fig 3B. Dashed grey line indicates DMSO average and dashed red line indicates 0.1nM colchicine average from Fig S3E; lead candidate Hit 2 is shown in red; n=3, mean±SEM.

**Figure S4. Hit 2 does not affect *ATXN* gene expression in control fibroblasts or rescue disease hallmarks in DM1 myotubes**

**A, B** Hit 2 does not affect expression of *ATXN1* in two out of three control fibroblast lines or of *ATXN3* in three control fibroblast lines at doses which reduce expression of equivalent expansion alleles (see Fig 5B-D); n=3, mean±SEM, **A** unpaired two-tailed t-test, **B** one-way ANOVA with Tukey’s multiple comparisons test, ns – not significant, * *P*<0.05.

**C** Three-day treatment of 1µM Hit 2 increases expression of *DMPK* in DM1 patient derived myotubes; n=3, mean±SEM, one-way ANOVA with Tukey’s multiple comparisons test, * *P*<0.05.

**D-F** Hit 2 treatment does not affect splicing of hallmark skipped exon events in DM1 myotubes, n=3, mean±SD.

**Figure S5. Mass spectrometry validation of in house synthesized Hit 2**

**A, B** Mass spectrometry of Hit 2 sourced from the NCI DTP (**A**) and in house synthesized Hit 2 (**B**). Top panels show total ion chromatograms and bottom panels show molecular ion peaks at 293.1729 (**A**) and 293.1979 (**B**).

**Figure S6. Widespread dysregulation of alternative splicing in *Atxn1^154Q/2Q^* SCA1 mouse model cerebellum**

**A** Percentage of significantly misspliced skipped exon (SE), retained intron (RI), mutually exclusive exons (MXE), alternative 5’ splice site (A5SS) and alternative 3’ splice site (A3SS) events as a proportion of total splicing events in DMSO injected SCA1 vs WT mice, number of each event shown on bar, FDR<0.1, ΔPSI>10%.

**B** Percentage of exon inclusion (positive) or exclusion (negative) for significantly alternatively spliced SE events, FDR<0.1, ΔPSI>10%, mean ± SD.

**C** Enrichment of summary gene ontology terms identified using Metascape for the 495 SE events dysregulated in DMSO injected SCA1 mice compared to DMSO injected WT mice.

**D, E** Violin plots of RNA-Seq data showing increased inclusion of *Trpc3* exon 9 (**D**) and increased exclusion of *Anks1b* exon 5 (**E**) in SCA1 mice; FDR<0.1, ΔPSI>10%, line indicates median.

**F** RT-PCR analysis of *Cacna2d2* exon 23 in cerebellum from presymptomatic (6 week, n=5), early symptomatic (12 week, n=4) and mild symptomatic (23 week, n=3) *Atxn1^154Q/2Q^* SCA1 mice compared to WT; ns – not significant, *** *P*<0.001, unpaired two-tailed t-test, mean±SEM.

**Supplemental Table 1. RNA Sequencing datasets**

