## supplemental figures for "CAG repeat-selective compounds reduce abundance of expanded CAG RNAs in patient cell and murine models of SCAs"

Figure S1

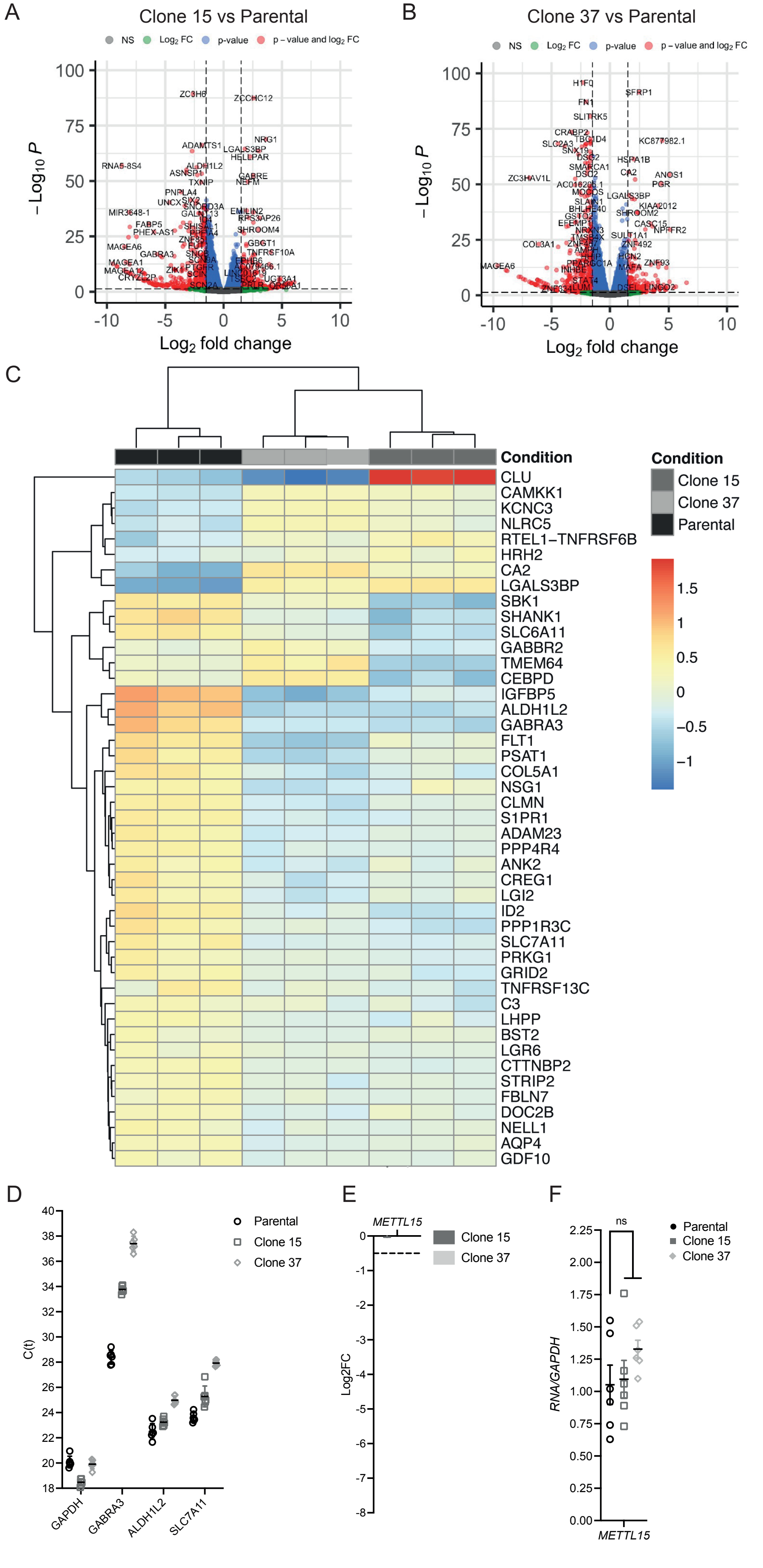
A
Clone 15 vs Parental
NS
Log2 FC
p-value
p – value and log2 FC
– Log10 P
Log2 fold change
B
Clone 37 vs Parental
C
Condition
Clone 15
Clone 37
Parental
CLU
CAMKK1
KCNC3
NLRC5
RTEL1–TNFRSF6B
HRH2
CA2
LGALS3BP
SBK1
SHANK1
SLC6A11
GABBR2
TMEM64
CEBPD
IGFBP5
ALDH1L2
GABRA3
FLT1
PSAT1
COL5A1
NSG1
CLMN
S1PR1
ADAM23
PPP4R4
ANK2
CREG1
LGI2
ID2
PPP1R3C
SLC7A11
PRKG1
GRID2
TNFRSF13C
C3
LHPP
BST2
LGR6
CTTNBP2
STRIP2
FBLN7
DOC2B
NELL1
AQP4
GDF10
1.5
1
0.5
0
–0.5
–1
D
C(t)
Parental
Clone 15
Clone 37
GAPDH
GABRA3
ALDH1L2
SLC7A11
E
METTL15
Log2FC
Clone 15
Clone 37
F
ns
Parental
Clone 15
Clone 37
RNA/GAPDH
METTL15

Figure S2

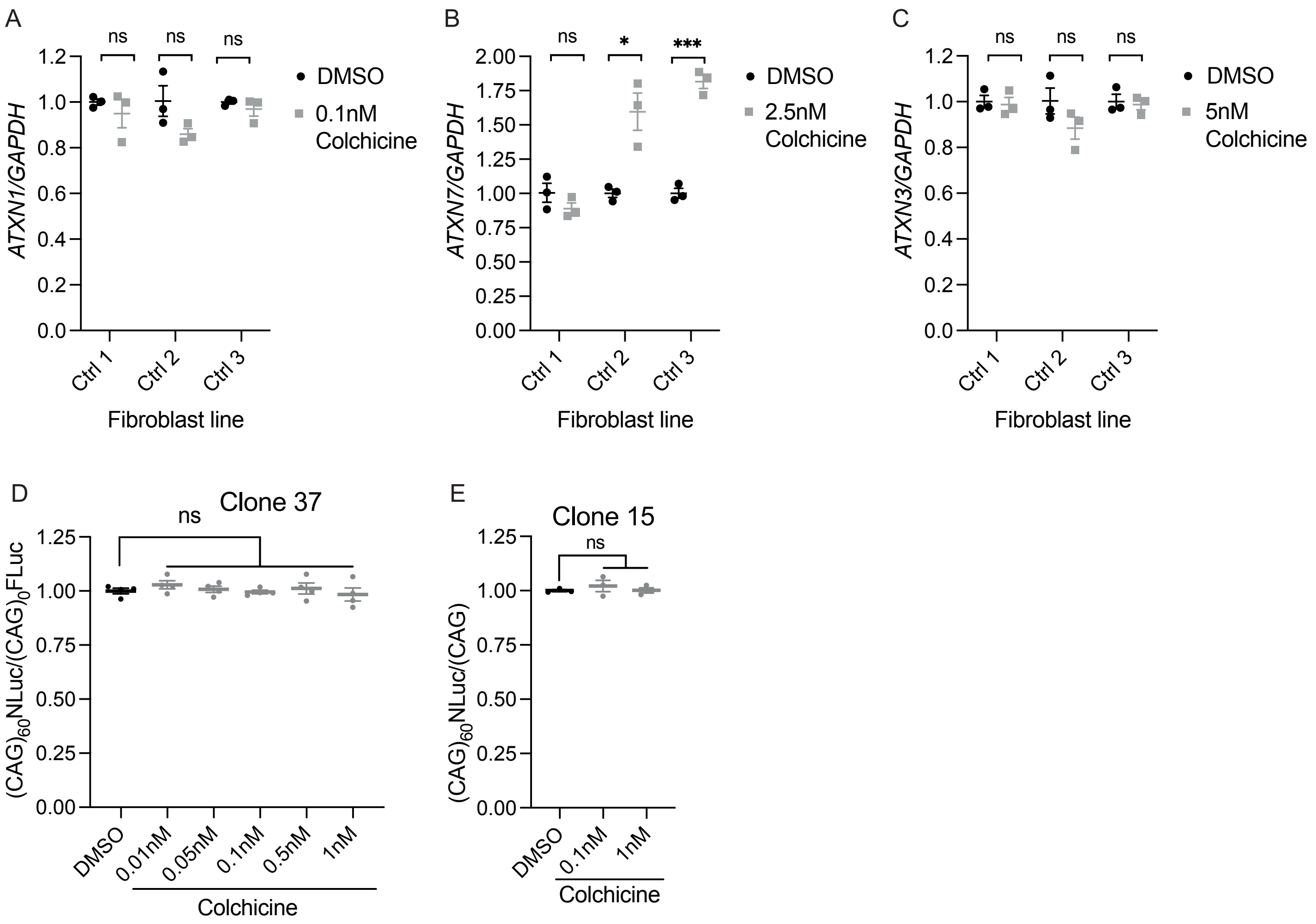

A
ns
ns
ns
1.2
1.0
0.8
0.6
0.4
0.2
0.0
ATXN1/GAPDH
Ctrl 1
Ctrl 2
Ctrl 3
Fibroblast line
DMSO
0.1nM Colchicine
B
ns
*
***
2.00
1.75
1.50
1.25
1.00
0.75
0.50
0.25
0.00
ATXN7/GAPDH
Ctrl 1
Ctrl 2
Ctrl 3
Fibroblast line
DMSO
2.5nM Colchicine
C
ns
ns
ns
1.2
1.0
0.8
0.6
0.4
0.2
0.0
ATXN3/GAPDH
Ctrl 1
Ctrl 2
Ctrl 3
Fibroblast line
DMSO
5nM Colchicine
D
Clone 37
ns
1.25
1.00
0.75
0.50
0.25
0.00
(CAG)60NLuc/(CAG)0FLuc
DMSO
0.01nM
0.05nM
0.1nM
0.5nM
1nM
Colchicine
E
Clone 15
ns
1.25
1.00
0.75
0.50
0.25
0.00
(CAG)60NLuc/(CAG)
DMSO
0.1nM
1nM
Colchicine

Figure S3

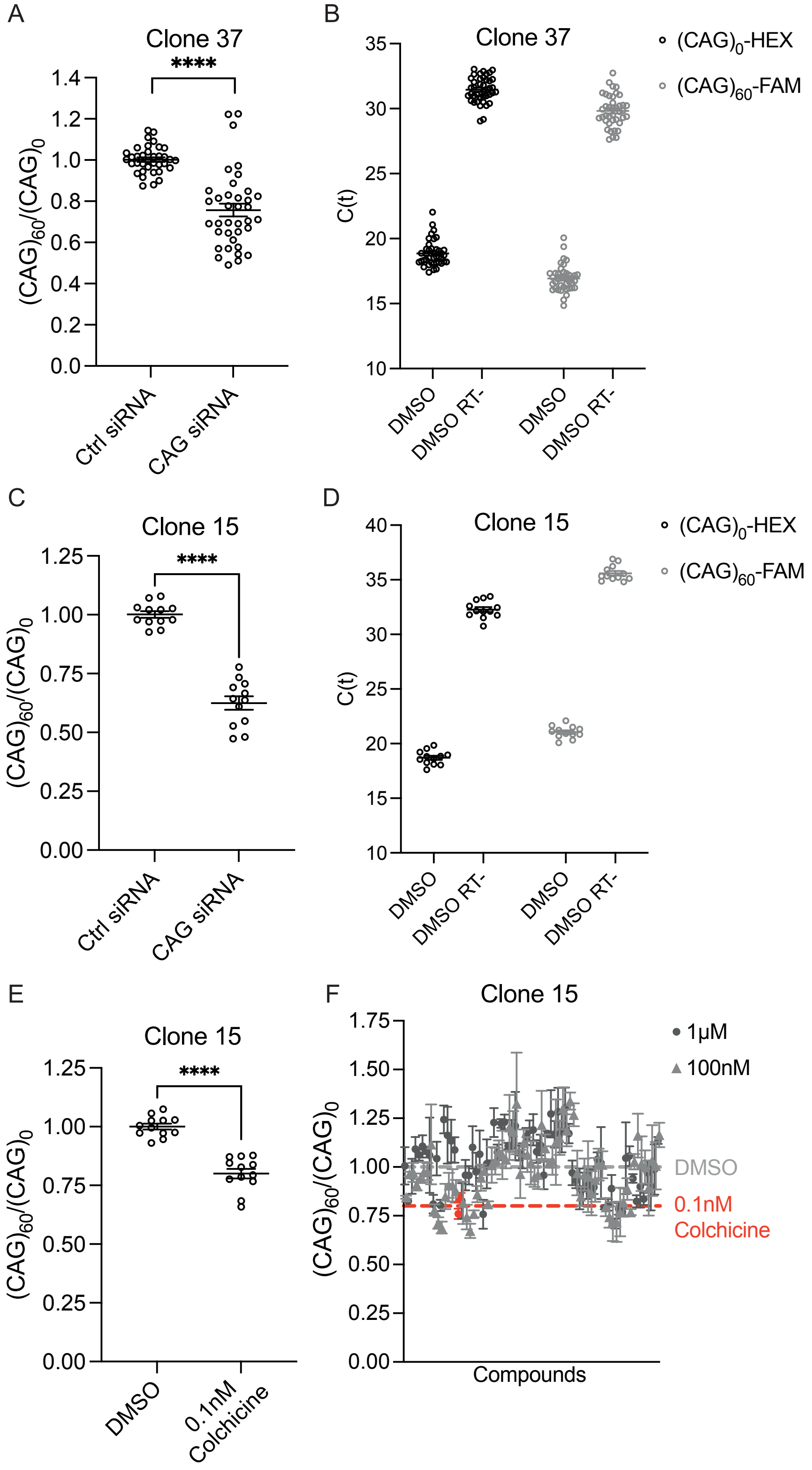

A
Clone 37
****
(CAG)60/(CAG)0
Ctrl siRNA
CAG siRNA
B
Clone 37
(CAG)0-HEX
(CAG)60-FAM
C(t)
DMSO
DMSO RT-
C
Clone 15
****
(CAG)60/(CAG)0
Ctrl siRNA
CAG siRNA
D
Clone 15
(CAG)0-HEX
(CAG)60-FAM
C(t)
DMSO
DMSO RT-
E
Clone 15
****
(CAG)60/(CAG)0
DMSO
0.1nM Colchicine
F
Clone 15
1μM
100nM
DMSO
0.1nM Colchicine
(CAG)60/(CAG)0
Compounds

Figure S4

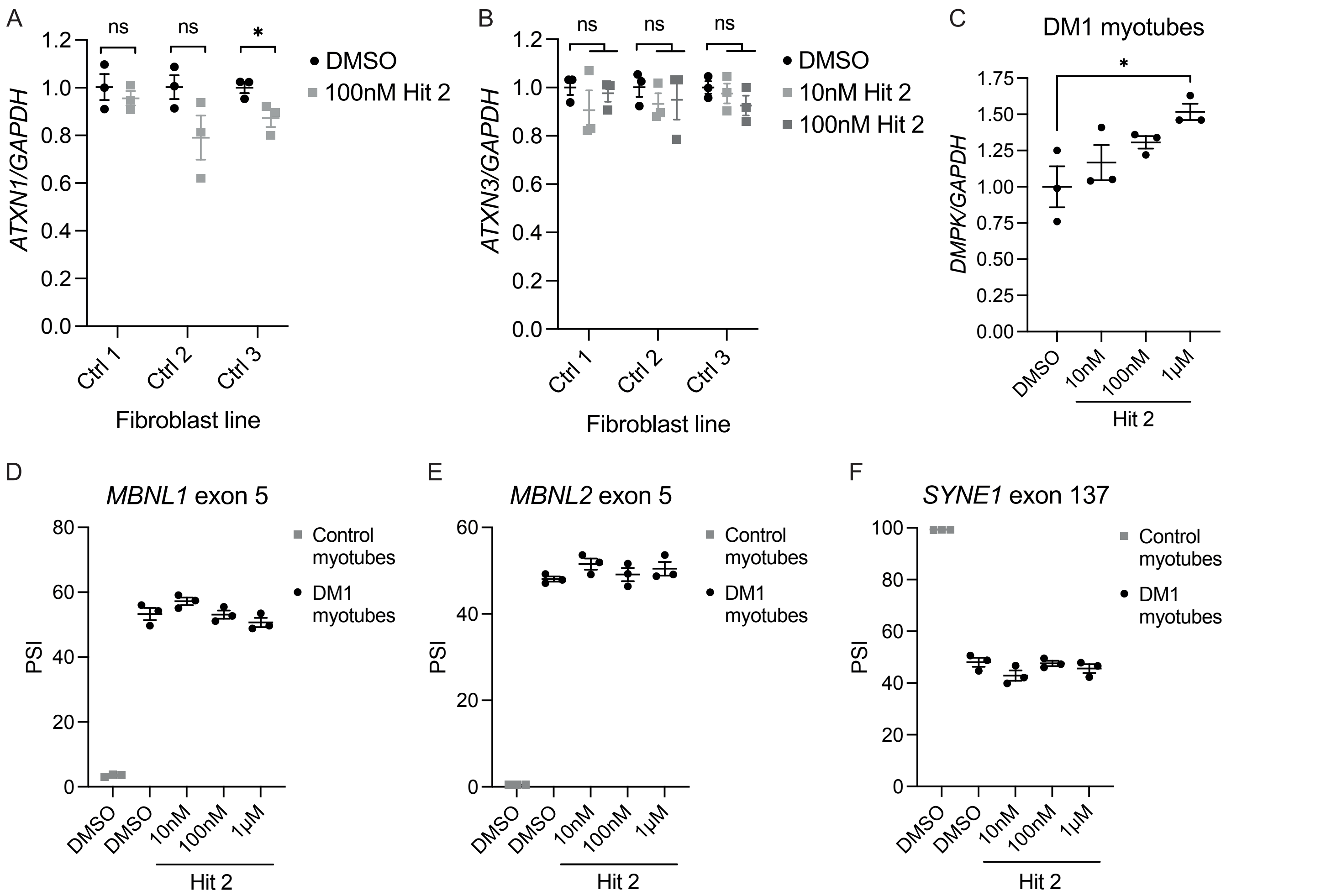
A
ns
ns
*
DMSO
100nM Hit 2
ATXN1/GAPDH
1.2
1.0
0.8
0.6
0.4
0.2
0.0
Ctrl 1
Ctrl 2
Ctrl 3
Fibroblast line
B
ns
ns
ns
DMSO
10nM Hit 2
100nM Hit 2
ATXN3/GAPDH
Ctrl 1
Ctrl 2
Ctrl 3
Fibroblast line
C
DM1 myotubes
*
DMPK/GAPDH
1.75
1.50
1.25
1.00
0.75
0.50
0.25
0.00
DMSO
10nM
100nM
1μM
Hit 2
D
MBNL1 exon 5
PSI
Control myotubes
DM1 myotubes
DMSO
DMSO
10nM
100nM
1μM
Hit 2
E
MBNL2 exon 5
PSI
Control myotubes
DM1 myotubes
Hit 2
F
SYNE1 exon 137
PSI
Control myotubes
DM1 myotubes
Hit 2

Figure S5

A

NCI-DTP Hit 2

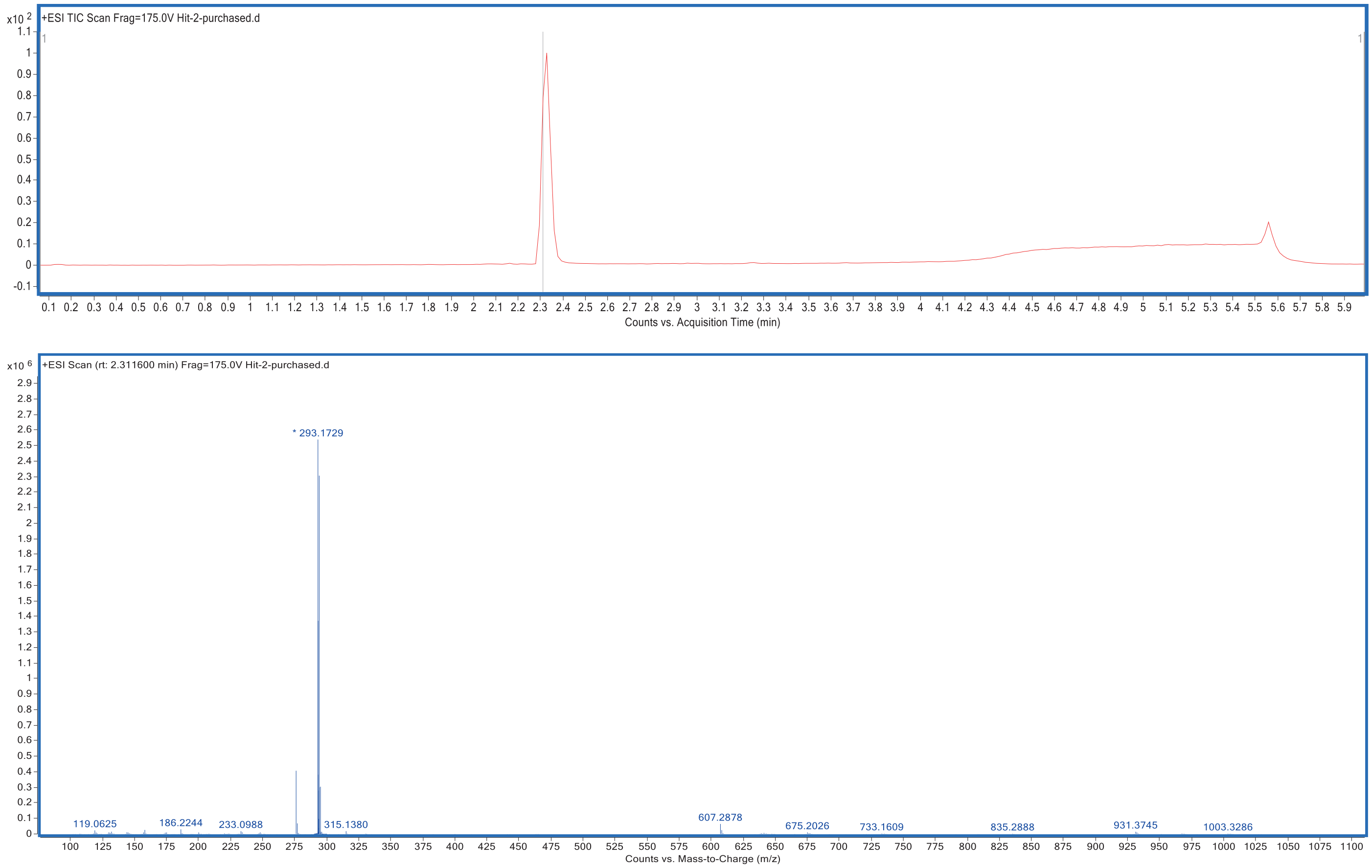


B

In-house synthesized Hit 2

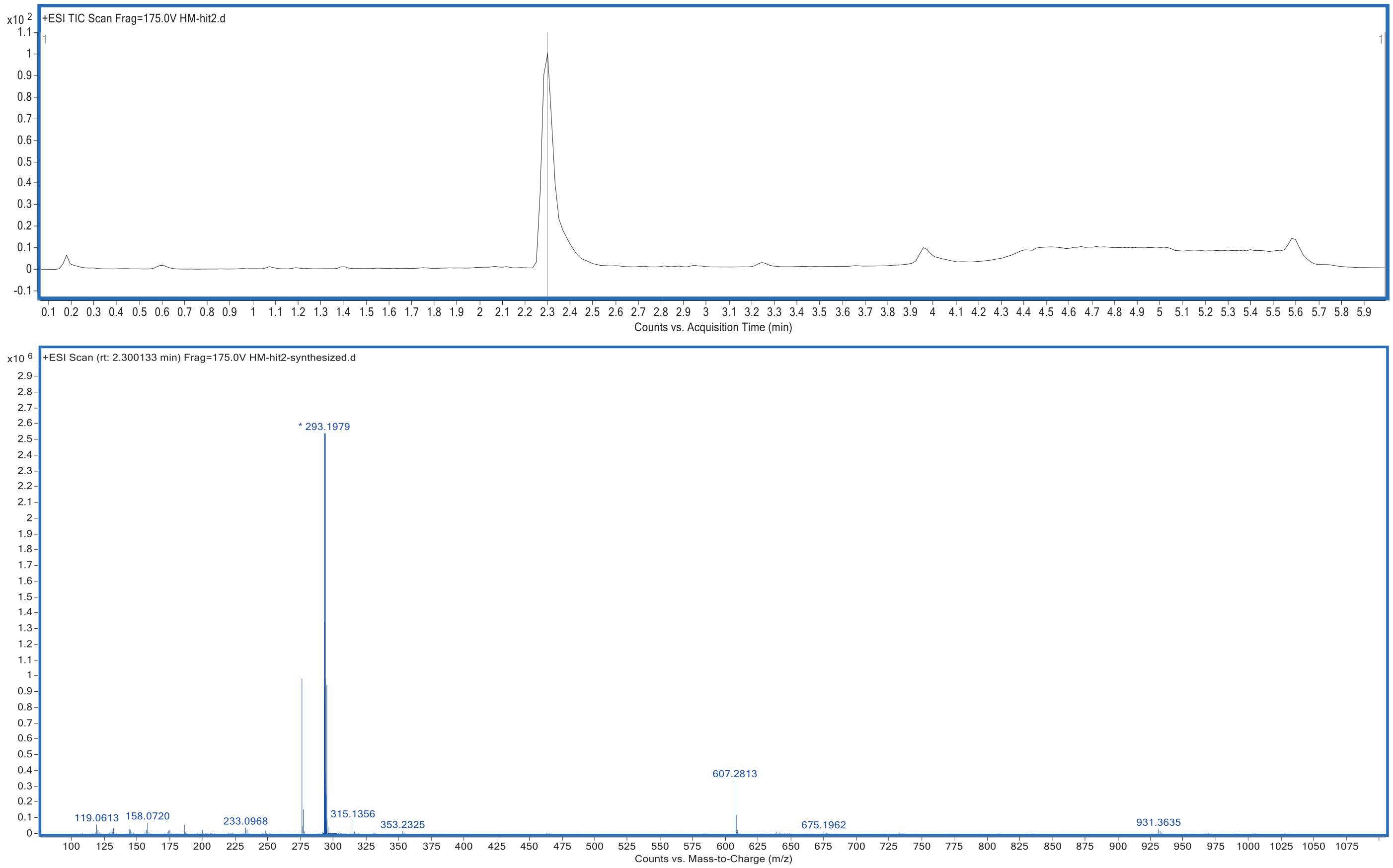

Figure S6

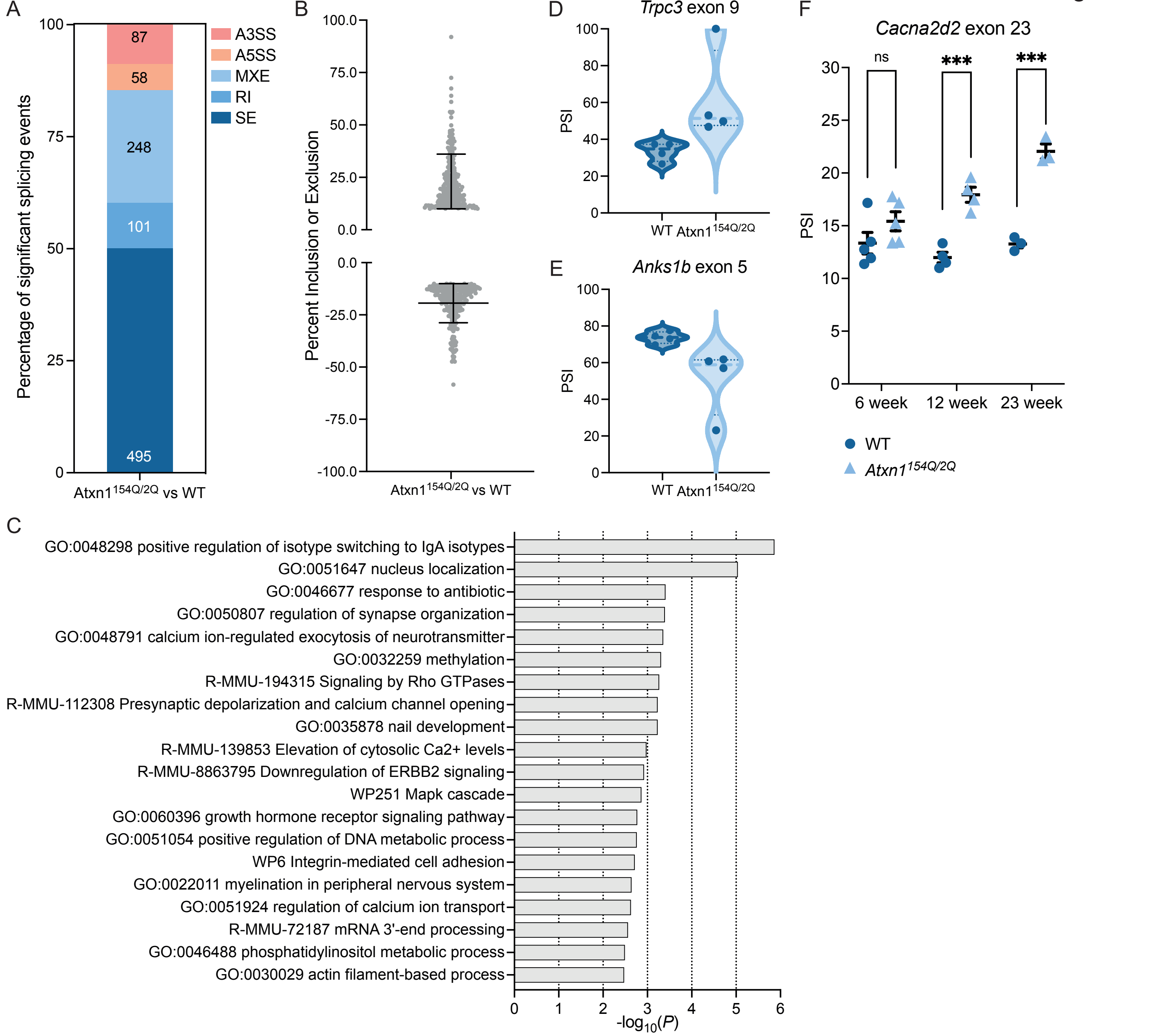

A
Percentage of significant splicing events
87
58
248
101
495
A3SS
A5SS
MXE
RI
SE
Atxn1154Q/2Q vs WT
B
Percent Inclusion or Exclusion
Atxn1154Q/2Q vs WT
D
Trpc3 exon 9
PSI
WT Atxn1154Q/2Q
E
Anks1b exon 5
PSI
WT Atxn1154Q/2Q
F
Cacna2d2 exon 23
PSI
ns
***
***
6 week
12 week
23 week
WT
Atxn1154Q/2Q
C
GO:0048298 positive regulation of isotype switching to IgA isotypes
GO:0051647 nucleus localization
GO:0046677 response to antibiotic
GO:0050807 regulation of synapse organization
GO:0048791 calcium ion-regulated exocytosis of neurotransmitter
GO:0032259 methylation
R-MMU-194315 Signaling by Rho GTPases
R-MMU-112308 Presynaptic depolarization and calcium channel opening
GO:0035878 nail development
R-MMU-139853 Elevation of cytosolic Ca2+ levels
R-MMU-8863795 Downregulation of ERBB2 signaling
WP251 Mapk cascade
GO:0060396 growth hormone receptor signaling pathway
GO:0051054 positive regulation of DNA metabolic process
WP6 Integrin-mediated cell adhesion
GO:0022011 myelination in peripheral nervous system
GO:0051924 regulation of calcium ion transport
R-MMU-72187 mRNA 3'-end processing
GO:0046488 phosphatidylinositol metabolic process
GO:0030029 actin filament-based process
-log10(P)

Table S1

| PRJNA1049475 | ID | Sex | Genotype | Treatment | RIN score | Total read-depth | Downsampled total read-depth |
|---|---|---|---|---|---|---|---|
| GSE249556 | 413 | M | Wt | DMSO | 8 | 147668268 | NA |
| | 425 | F | Wt | DMSO | 10 | 226178630 | 130000000 |
| | 426 | F | Wt | DMSO | 10 | 217174296 | 130000000 |
| | 431 | M | Wt | DMSO | 9.2 | 122989072 | NA |
| | 409 | M | Het | DMSO | 6.4 | 212785844 | 130000000 |
| | 411 | M | Het | DMSO | 7 | 88591702 | NA |
| | 420 | M | Het | DMSO | 9.8 | 87317374 | NA |
| | 429 | M | Het | DMSO | 8.3 | 129594098 | NA |
| | 410 | M | Het | Hit 2 | 8.5 | 86298122 | |
| | 421 | M | Het | Hit 2 | 7.5 | 108777144 | NA |
| | 422 | M | Het | Hit 2 | 9.8 | 239793282 | 140000000 |
| | 430 | M | Het | Hit 2 | 9.5 | 11221174 | NA |
| GSE249555 | P1_Parental | NA | Parental | NA | NA | 39997046 | NA |
| | P2_Parental | NA | Parental | NA | NA | 46576068 | NA |
| | P3_Parental | NA | Parental | NA | NA | 49212254 | NA |
| | C1_Clone37 | NA | Clone 37 | NA | NA | 40939306 | NA |
| | C2_Clone37 | NA | Clone 37 | NA | NA | 40565810 | NA |
| | C3_Clone37 | NA | Clone 37 | NA | NA | 44406142 | NA |
| | C1_Clone15 | NA | Clone 15 | NA | 10 | 139405850 | NA |
| | C2_Clone15 | NA | Clone 15 | NA | 10 | 92028910 | NA |
| | C3_Clone15 | NA | Clone 15 | NA | 10 | 68553816 | NA |
